# Temporal occupancy distributions reveal multifaceted and heterogeneous effects of climatic variation on montane butterflies

**DOI:** 10.1101/2025.02.07.637160

**Authors:** Gbolahan A Reis, Matthew L Forister, Christopher A Halsch, Clare M Dittemore, Arthur M Shapiro, Zachariah Gompert

## Abstract

Climate change has substantially altered the phenology of many organisms, with profound implications for biodiversity and species interactions. However, phenological studies of insects have often focused on shifts over time, without explicitly examining how climate drives these changes. In addition, the metrics used to measure changes in phenology often capture only limited information about the flight period. In this study, we analyzed multidecadal observational and climate data using a hierarchical Bayesian framework to model the annual probability of occurrence distributions for 135 butterfly species across five montane sites along an elevational gradient. Our analysis used polynomial models that account for shifts in abundance and the timing and length of flight periods to investigate the effect of climate on butterfly phenology. We found that spring maximum and minimum temperatures, as well as winter precipitation, are important predictors of butterfly phenology. High winter precipitation delayed phenology at high-elevation sites where substantial snowfall occurs, while increased spring maximum temperatures generally advanced phenology across all elevations. Even modest increases in spring minimum temperatures caused substantial shift in phenology. We documented variability in the effect of climate on phenology across sites, among species within a site and among populations of the same species across different sites with variability among species within a site being especially pronounced. We also found that climate influences different aspects of the flight period differently (e.g., timing versus duration), underscoring the need to move beyond single metrics such as day of first flight. These findings highlight the importance of examining the entire flight period and considering the interplay of species-specific traits to improve predictions of how climate change impacts phenology. Such approaches might be essential for designing more targeted and effective conservation strategies in response to ongoing climate change.

## 1 INTRODUCTION

Climate change is reshaping ecosystems around the world, with numerous studies documenting substantial impacts on the ecology of organisms (Halsch et al. 2021, Inouye 2022, Parmesan 2006). Among the most evident biological responses to climate change are phenological shifts, with species experiencing changes in the timing of critical life history events such as migration, flowering, and flight periods (Cleland et al. 2007, Forister et al. 2018, Newson et al. 2016, Parmesan 2006 2007). These changes in phenology have implications for biodiversity, ecosystem services, and species interactions (Kharouba et al. 2018, Miller-Rushing et al. 2010, Renner and Zohner 2018, Thackeray et al. 2010). Consequently, understanding and predicting these shifts has become a focal point for ecologists, especially in the context of ongoing global climate change.

In ectothermic organisms, such as insects, phenological responses are particularly pronounced because their metabolic and developmental rates are directly influenced by external temperature and precipitation patterns (Cohen et al. 2018, Parmesan 2006, Deutsch et al. 2008). Earlier flight periods have been widely documented in regions experiencing increased temperatures (Forrest 2016), with evidence that these shifts depend on both the magnitude of climate warming and species-specific sensitivity to environmental changes (Chmura et al. 2019).

Butterflies serve as an excellent system for investigating shifts in flight periods. Studies on the impacts of climate warming on butterfly flight periods have mainly focused on long-term trends, often without directly examining how annual climate conditions affect yearly flight periods.. These studies have documented earlier flight periods, increased voltinism, and extended flight periods (Colom et al. 2021, Forister and Shapiro 2003, Gutiérrez and Wilson 2021, Habel et al. 2024). However, the magnitude and direction of these shifts are not uniform across species or populations (Diez et al. 2012), and this variability is influenced by differences in ecological and life history traits (Diamond et al. 2011, Forrest 2016, Zografou et al. 2021). Additionally, local adaptation and phenotypic plasticity play critical roles in shaping responses to climate, leading to diverse patterns across habitats and geographic regions (Colom et al. 2021, Gutiérrez and Wilson 2021, Habel et al. 2024). Shifts in the flight period of butterflies can have cascading effects on ecological interactions, potentially leading to mismatches between species that rely on synchronized timing for mutualistic relationships, such as pollinators and flowering plants. While mismatches are expected to negatively impact fitness, evidence for such mismatches in ecological interactions is currently weak or mixed (Kharouba and Wolkovich 2023).

Traditionally, shifts in flight periods have been quantified using simple metrics like the first, mean, or last day of flight (Falt ý nek Fric et al. 2020, Parmesan 2007, Roy and Sparks 2000, Thackeray et al. 2016). However, these metrics are limited, as they are influenced by factors such as species abundance, period of observation, and sampling effort (Van Strien et al. 2008, Inouye et al. 2019, Pearse et al. 2017, Miller-Rushing et al. 2008). They also fail to capture intra-annual variability or the uncertainty surrounding species occurrence during the flight period (i.e. the occurrence of individuals on other potentially unmonitored days within the flight season), thus representing only a fraction of the full flight period. Consequently, these metrics provide an incomplete picture of how consistently and to what extent flight periods are shifting over time. Focusing on these limited metrics overlooks the effect of climate on other aspects of the flight period, which could substantially impact species interactions and broader ecological dynamics (Forrest and Miller-Rushing 2010, Macphie et al. 2023).

Furthermore, climate-driven changes in one aspect of the flight period might not accurately represent effects on other facets of the flight period including the length of the flight window or the date of last flight. For example, while rising temperatures might lead to earlier emergence, this does not necessarily imply an extended overall flight season. This discrepancy introduces challenges in our understanding of mismatches, particularly as meta-analyses and reviews often compare studies that measure different and distinct aspects of the flight period. Such differences complicate cross-study comparisons and may obscure broader patterns (Cohen et al. 2018). To improve predictions of species’ responses to climate change and the associated ecological implications, a more comprehensive understanding of climate’s impact on flight period is essential.

This study uses a comprehensive long-term butterfly observational dataset, encompassing 135 species across various elevations and habitat types in the Sierra Nevada mountains, to understand changes in flight periods by modeling annual occurrence distributions, that is, the probability of occurrence or detection across the flight period for each species at each site in each year (Figure 1). Montane habitats are particularly vulnerable to climate change, with temperatures rising more rapidly at higher elevations (Ohmura 2012). Previous studies at these sites have focused on the effect of climate on abundance (as captured by average probability of occurrence for a butterfly species in a year) rather than phenology. These studies showed that the effects of climate on butterfly abundance vary substantially among different sites, highlighting the heterogeneous nature of climate responses (Nice et al. 2019). Some work has been done with these data on phenology but only in a limited and focused way. For example, drought events were shown to compress flight windows in univoltine species and reduce species richness, indicating the potential for climate extremes to significantly alter flight periods (Forister et al. 2018).

**FIGURE 1.**
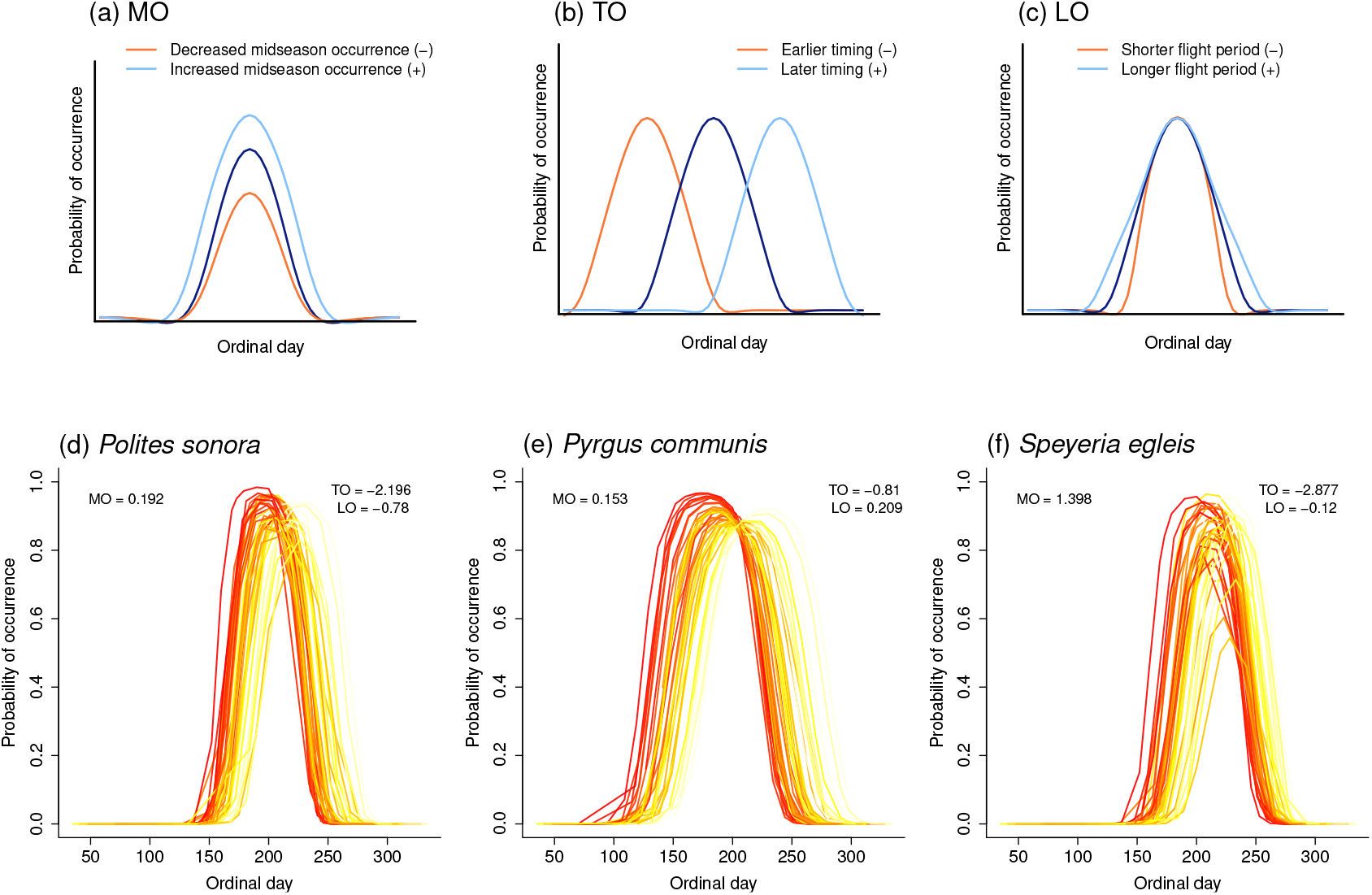
Effects of climate on the annual occurrence distribution of butterfly species. Panel (a) indicates the effect of climate on mid-season occurrence (MO), with negative effects denoting a decreased probability of occurrence and positive effects indicating an increased probability of occurrence across the flight period. Panel (b) shows the impact of climate on the timing of occurrence (TO) (interaction between climate variables and ordinal day), with negative effects indicating a shift to an earlier flight period and positive effects suggesting a shift to a later flight period. Panel presents the effect of climate on the length of occurrence (LO) (interaction between climate variables and ordinal day^2^), with negative values indicating a shorter flight period and positive values indicating a longer flight period. Panels (d), (e) and (f) show estimates of interannual variation in flight periods for three representative species *Polites sonora, Pyrgus communis* and *Speyeria egleis* at Donner Pass from our climate model. Each line represents the probability of occurrence on each day across a year, with colors indicating average spring maximum temperatures for that year (darker colors indicate higher spring maximum temperatures). Thus the effects of spring maximum temperature on MO, TO and LO for each species are shown in each plot.

We combine this butterfly dataset with climate data and use hierarchical Bayesian generalized linear models to assess the effects of climate on both intra-annual and inter-annual variation in butterfly occurrence distributions. Our analysis allows us to model the effects of climate on the probability of occurrence for butterflies across the season (i.e., the occurrence distribution) in three distinct ways. Specifically, our models include climatic effects on: (i) the mid-season probability of occurrence (MO), which is an intercept term that specifically parameterizes the probability of occurrence on ordinal day 0 (after centering, hence the term mid-season) but affects the probability of occurrence throughout the entire flight season; past work has shown that the probability of occurrence throughout the flight season is a good metric of abundance and thus the effect of climate on MO captures the effect of climate on abundance (Casner et al. 2014, Halsch et al. 2024); (ii) the timing of the flight period (i.e., the timing of occurrence or TO); (iii) the duration of the flight season, which we refer to as the length of occurrence (LO) (Figure 1). This approach accounts for species- and site-specific responses, providing a more nuanced understanding of how climatic effects could differ by species and sites. Our study first focuses on examining how each aspect of the occurrence distribution has changed over time without directly modeling climatic factors. We then investigate how inter-annual variation in climate affects inter-annual variation in the occurrence distribution, as captured by the effect of climate on MO, TO, and LO. We then explore how climate effects vary across sites, among species within sites, and among populations of the same species across sites. Finally, we ask whether climatic variation has similar effects on different aspects of the flight period, that is on MO, TO and LO. Our findings emphasize the importance of considering the entire flight period, rather than distinct aspects alone, to improve our understanding of how species respond to climate change.

## 2 MATERIALS AND METHODS

### 2.1 Butterfly Dataset

The butterfly dataset used in this study is part of a long-term monitoring program that was begun in the early 1970s by one of us (AMS), encompassing 10 sites across Northern California. Since 2018, monitoring at the highest elevation sites has been conducted by three of us (MLF, CAH, CMD) using the same methodology. Here, we focused on observations from five sites–Castle Peak (CP), Donner Pass (DP), Lang Crossing (LC), Sierra Valley (SV) and Washington (WA)–in the Sierra Nevada mountains, spanning an elevational gradient from 800m to 2800m (Figure 2 a, b). These sites exhibit diverse climate conditions (Figure 2 c, d, e) and habitat types. Surveys at these sites are conducted every other week during the butterfly flight season on days conducive to butterfly activity. The number of years that each site has been monitored varies, some starting as early as 1973 (DP), while others began later (SV in 1982, WA in 1988). The dataset for sites which were monitored before 1985 was truncated so that the earliest observations were in 1985. During each survey visit, observers followed a fixed, permanent route and recorded the presence or absence of all butterfly species. Detailed information on each site and the corresponding survey routes can be found in Shapiro (2024). For this study, at each site, we focused on butterfly species observed at least ten times throughout the study years, excluding strays and infrequent species. This selection resulted in 64 species at CP, 88 species at DP, 83 species at LC, 70 species at SV, and 80 species at WA. A total of 104 species were found at multiple sites (i.e., at least two sites), with 26 species shared across all five sites, contributing to an overall total of 135 unique species observed across the study sites (Figure A1).

**FIGURE 2.**
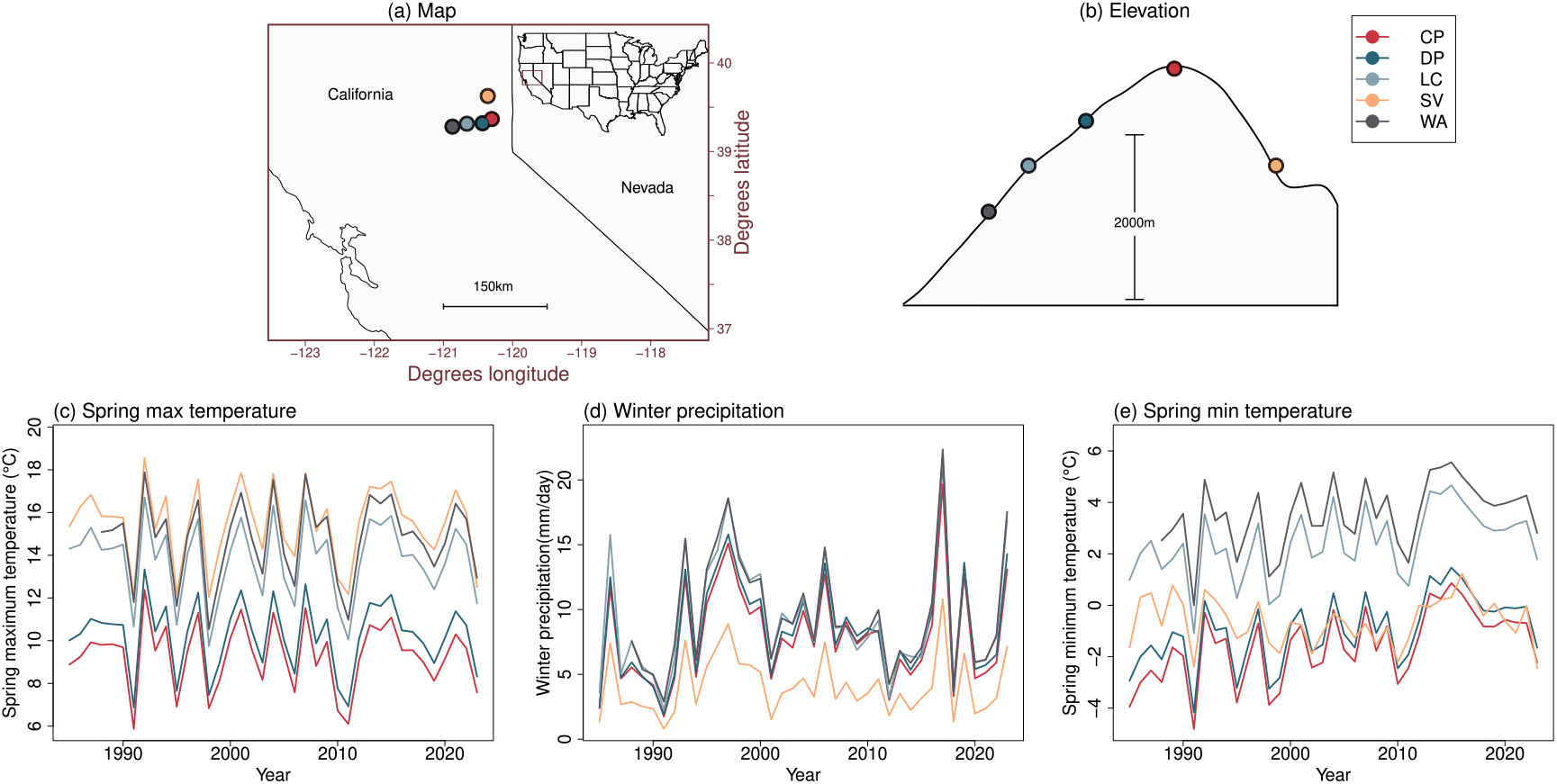
Site locations and annual climatic variation. Panel (a) is a map showing the five study sites in the Sierra Nevada with an inset of a USA map with the focal area enclosed in a box. Panel (b) shows the elevation at each site. Panel (c), (d) and (e) display inter-annual variation in average spring maximum temperature (c), average winter precipitation (d) and average spring minimum temperature (e) across sites from 1985 to 2023. Site abbreviations are as follows: CP (Castle Peak), DP (Donner Pass), LC (Lang Crossing), SV (Sierra Valley) and WA (Washington).

### 2.2 Butterfly Modeling

To quantify shifts in butterfly flight periods, we modeled their annual occurrence distributions using a hierarchical Bayesian framework. Butterfly occurrence on a given day was modeled as a Bernoulli random variable, where the presence of a butterfly on a specific ordinal day was classified as a success. We modeled the probability of occurrence for species i on ordinal day j in year k (denoted as P_(i, j, k)_) using a generalized linear model (Equations 1 and 2). Since the probability of occurrence of butterflies typically follows a parabolic pattern within a year–first increasing and then decreasing–we included a quadratic polynomial for ordinal day (i.e., including both ordinal day and ordinal day^2^ terms) to capture these dynamics. First, we fit a baseline time model, and then considered various suites of climatic variables. For the time model, the design matrix (**X**) included ordinal day, ordinal day^2^, year, and the interactions between year and both ordinal day and ordinal day^2^. For the climate models, the design matrix (**X**) included climate variables, ordinal day, ordinal day^2^, year, and the interactions between each climatic variable and both ordinal day and ordinal day^2^.

We estimated species-specific intercepts (*α*_i_) and effects for explanatory variables (*β*_i_)–all of which were Z-standardized–within the matrix (**X**) (Equation 2). The hierarchical mean (*μ*) and hierarchical standard deviation (*σ*) denote the mean and variability of the species-specific effects of each explanatory variable at each site (Equation 3). The model specification was as follows:

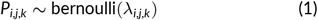

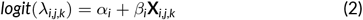

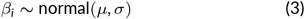

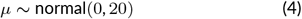

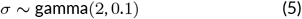

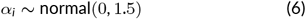

The main effect for each explanatory variable (climate variable or year) represents the direct effect of that variable on MO, where negative and positive effects indicate decreased and increased overall probabilities of occurrence (and thus abundance), respectively. The interaction effect between each explanatory variable and ordinal day represents the effect of that variable on TO, with negative effects indicating a shift to an earlier flight season and positive effects indicating a shift to a later season. These shifts apply to the entire probability of occurrence distribution. The interaction effect between each explanatory variable and ordinal day^2^ represents the effect of that variable on LO, where negative values indicate a shorter flight season, and positive values a longer season (this assumes a convex function for the probability of occurrence across the season) (Figure 1 a, b, c).

These curves modeled using the biweekly presence and absence observations for each species effectively describe the timing, length and mid-season occurrence of butterfly species, capturing changes in the probability of occurrence across the flight period (Figure 1). Separate models were fitted for each site. Our hierarchical approach enabled us to estimate the effects of each explanatory variable at two levels: for individual species within a site (captured by the *β* coefficients) and across all species at a site (captured by the parameter *μ*), thus capturing both species-specific and site-level effects and variability (with the latter captured by the parameter *σ*). To maintain consistent interpretation of the regression coefficients, we excluded species whose peak probability of occurrence fell at the beginning and end of the flight season, that is where the probability of occurrence distribution was convex rather than concave (this is expected for some species that overwinter as adults). This included five species at CP, one species at LC and two species at SV.

All analysis were conducted in R (Version 4.2.2). We used relatively vague priors for all parameters estimated in our analysis (Equations 3, 4, 5, 6). Posteriors were sampled using Hamiltonian Monte Carlo in Stan via the RStan package (Version 2.21.8). Four chains were run, each with 4,000 iterations, and the first 2,000 iterations were considered as the burn-in period. Model convergence was assessed through Gelman–Rubin diagnostics and trace plots. Models were compared using Pareto-smoothed importance sampling and posterior predictive checks using the loo package (Version 2.5.1). The posterior distribution of each estimated parameter was summarized by calculating its median and 95% credible interval (CrI).

### 2.3 Temporal trends in flight period

To examine overall temporal trends in butterfly abundance and phenology, we fitted a baseline time model using the year (effect of year on MO), ordinal day of occurrence within that year, ordinal day^2^, interaction between year and ordinal day (effect of year on TO) and interaction between year and ordinal day^2^ (effect of year on LO) as explanatory variables. This initial model provided a framework to capture time-related changes in the period of flight, serving as a baseline to assess temporal trends. However, while time reflects overall patterns, it is not a causal factor. Thus, we further examined specific climatic factors that may underlie these trends and also explain the inter-annual variation in flight period.

### 2.4 Effect of climate on the annual flight period

To investigate the effects of climate on annual flight period of butterflies, we obtained daily climate data for each site, including minimum temperature, maximum temperature, precipitation, and snow water equivalent, from Daymet daily surface weather and climatological summaries, corresponding to the Daymet grid pixel (1×1 km spatial resolution) (Thornton et al. 2022). Minimum temperature, maximum temperature and precipitation were averaged at both annual and seasonal levels, with seasons defined as follows: fall summaries represented the average daily climate data from September to November of the previous year, winter summaries covered December of the previous year through February of the current year, and spring summaries included March to May of the current year. We also calculated the mean daily snow water equivalent (SWE) from September 1 of the previous year to May 31 of the current year. Using daily minimum and maximum temperature we estimated growing degree days (GDD) accumulated from January 1 to May 31 of the current year using methods described in Cayton et al. (2015) with a minimum and maximum threshold of 10°C and 30°C respectively. We considered eleven climatic variables, including the mean daily precipitation, daily minimum temperature, and daily maximum temperature for fall, winter, and spring, as well as GDD and SWE. We specifically focused on climate summaries from the previous fall, winter, and spring to understand how climate influences phenology as mediated through overwintering and early season developmental stages.

To identify the climatic variables that best explained variations in annual flight period, we employed a forward model selection approach (Table A1). We began by fitting eleven single-climate variable models, each incorporating a climatic variable as an additional variable to the explanatory variables within the baseline time model (this included effects of the variable on MO, TO and LO). We selected the model with the best predictive performance based on approximate leave-one-out cross-validation (Vehtari et al. 2017). Using the best single-climate variable model, we constructed ten two-climate variable models by adding a second climatic variable and assessed whether this addition improved model performance. If the model performance improved, we selected the best two-climate variable model and repeated the process to fit models with three climatic variables. This iterative process continued until the inclusion of an additional climatic variable no longer increased model predictive performance (Table A1). Model comparisons, including the baseline time model, were conducted by estimating the expected log pointwise predictive density (ELPD) via approximate leave-one-out cross-validation using the loo package (Version 2.5.1). The ELPD values were averaged across all sites for each model, and the model with the highest average ELPD was selected as the best model. All models incorporating climatic variables outperformed the baseline time model, suggesting that annual climate could account for additional variation in flight period. The model that included spring maximum temperature, spring minimum temperature, and winter precipitation as climatic variables best explained the flight period of butterflies on average across all sites (Table A1). Consequently, we used this model to investigate the effects of climate on the annual MO, TO, and LO of butterfly species.

To investigate variation in the effects of climate on the different model parameters describing the probability of occurrence distribution, we quantified variation in effects at three distinct levels. First, we assessed how the average effects of each climatic variable on MO, TO and LO varied across sites by calculating the standard deviation of the estimated mean effects (the *μ*s) across the five sites. Second, we evaluated the level of variation among species within sites on average by estimating the mean of all hierarchical standard deviations among species within each site (i.e., the mean of the estimated *σ* parameters from the model). Lastly, we examined variation among populations of species found in multiple sites on average. This was accomplished by estimating the standard deviation in effects of climate on populations of each of the 104 species present in multiple sites and averaging these estimates across species.

To determine whether differences in the average effects of climatic factors at each site were caused by differences in species compositions versus differences in how the same species were affected by climate, we examined the consistency of climate effects across species shared between each pair of compared sites. High correlations in the effects of climate between sites would suggest that differences in species composition drive differences in average effects among sites, offering insight into how community composition shapes site-level climate responses.

Additionally, we investigated whether the effects of climate on different facets of the occurrence distribution are associated, as well as how these correlations varied across sites. To do this, we estimated the correlation between the species-specific effect of each climatic variable on TO versus LO, TO versus MO and MO versus LO of species at each site.

Because climate variables are multifaceted, often correlated, and interact with each other, we also employed a multivariate approach using principal component analysis (PCA). This approach allowed us to simultaneously consider all climate variables and examine how their effects on different facets of the occurrence distribution were correlated. For each site, a PCA was performed using the species-specific effects of each climatic variable on each aspect of the flight period for species observed at that site. As part of this analysis, we examined whether the correlations between effects of climate on different facets of the occurrence distribution varied between univoltine and multivoltine species as such species are expected to be differently affected by climatic variation (Diamond et al. 2011).

Finally we examined whether the associations between the effects of climate on different aspects of the flight period show similar patterns across sites. To do this, we conducted an additional PCA that focused only on species found at all sites, ensuring that differences in species composition do not influence the results. For this, PCA was performed using species-specific effects of each climatic variable on each aspect of the flight period across all five sites.

## 3 RESULTS

### 3.1 Temporal trends in the occurrence distribution of butterfly species

We examined temporal changes in flight period as captured by the effect of year on MO, TO and LO parameters describing the probability of occurrence across the flight period. Despite species-specific variability, we observed a consistent decline in MO across all sites, indicating a general decline in the average probability of occurrence for butterflies over time (Figure 3 a). Additionally, species-level variation was evident in TO, with some species shifting toward earlier emergence, while others had a delayed flight season (Figure 3 b). Similar species-specific patterns were observed for LO, with varying trends in the length of the flight period across species (Figure 3 c).

**FIGURE 3.**
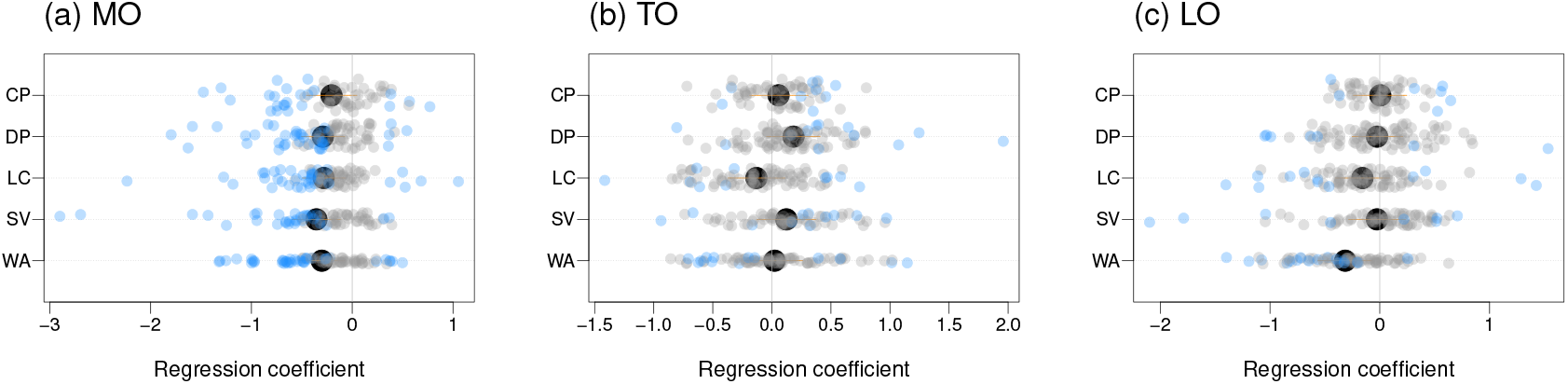
Temporal trends in butterfly flight period at each site. Each panel shows the effect of year on the (a) mid-season occurrence (MO), (b) timing of occurrence (TO) and (c) length of occurrence (LO) for each butterfly species at each site. In each panel, large black points with orange bars show the site-level response and 95% credible intervals (CrIs). The small points show population-specific responses; blue colored points denote credible population-level effects whereas grey colored points indicate cases where 95% CrIs for population effects include 0.

### 3.2 Effects of climate on the occurrence distribution of butterfly species

Models incorporating climatic variables outperformed the time model, suggesting that annual climate explains additional variation in annual occurrence distributions and thus in phenology. The model with the highest predictive power across sites included spring maximum temperature, spring minimum temperature, and winter precipitation as predictors, suggesting these climatic factors explain inter-annual variation in butterfly abundance and phenology (Figure A2 A35).

Overall, we did not detect credible site-level effects or consistent species-specific effects within species at each site for most climatic variables on MO, and thus on the overall probability of occurrence across the year, which is related to abundance (Figure 4 a, b, c). However, a subset of species exhibited credible effects of each climatic variable on MO at each of the five sites. Moreover, at CP, we found a credible negative site-level effect of spring maximum temperature (−0.47, 95% CrI: -0.72 to -0.23) and this was consistent across species with credible species-specific effects, suggesting that higher spring maximum temperatures reduced MO (Figure 4 a). Similarly, at DP, we found a credible negative site-level effect of spring minimum temperature (−0.38, 95% CrI: -0.58 to -0.18) and this was consistent across species with credible species-specific effects, suggesting that higher spring minimum temperatures reduced MO (Figure 4 c).

**FIGURE 4.**
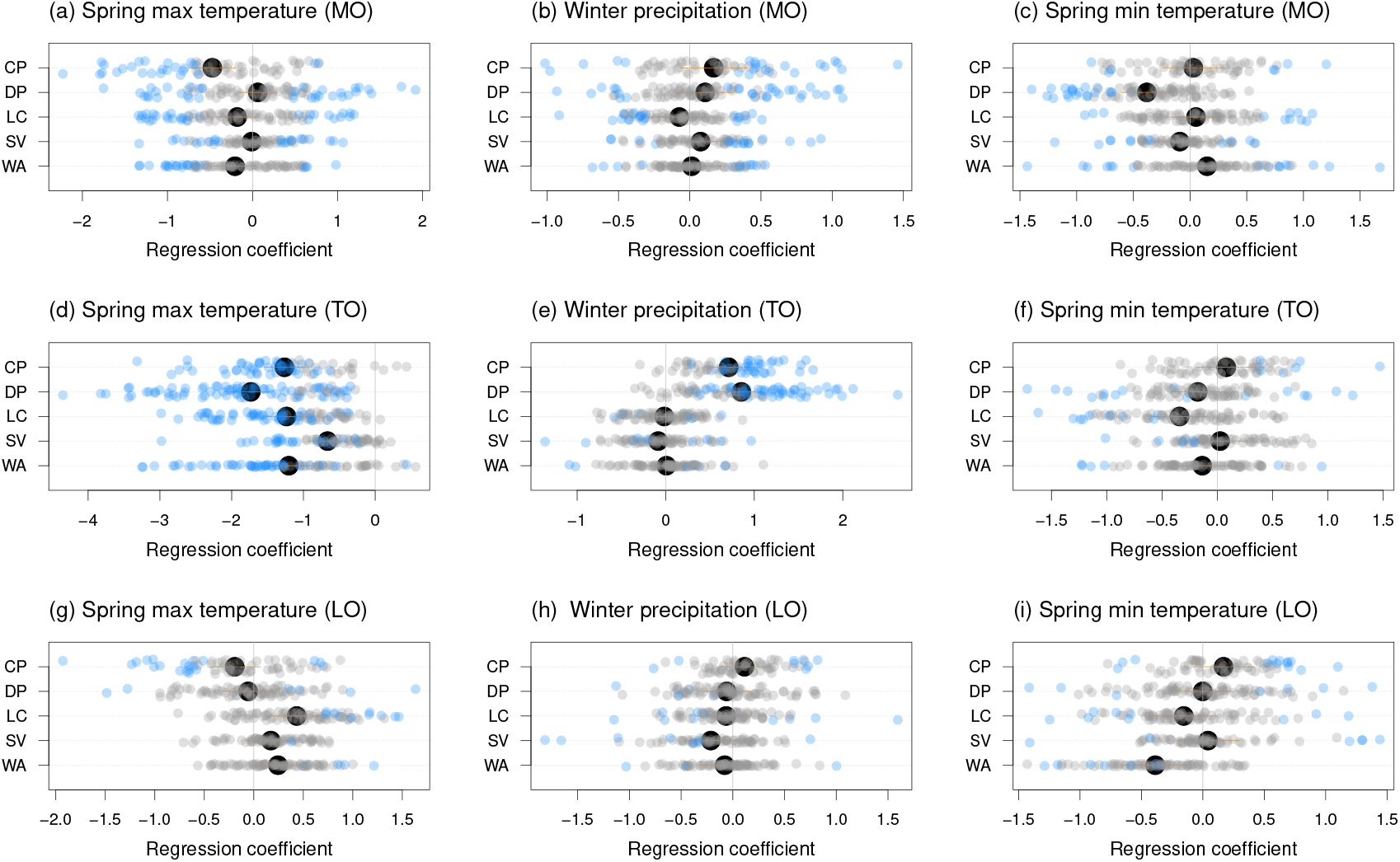
Effect of climate on butterfly flight period at each site. Results are shown for the effect of (a) spring maximum temperature on the mid-season occurrence (MO), (b) winter precipitation on the mid-season occurrence (MO), (c) spring minimum temperature on the mid-season occurrence (MO), (d) spring maximum temperature on the timing of occurrence (TO), (e) winter precipitation on the timing of occurrence (TO), (f) spring minimum temperature on the timing of occurrence (TO), (g) spring maximum temperature on length of occurrence (LO), (h) winter precipitation on length of occurrence (LO), (i) spring minimum temperature on length of occurrence (LO). In each panel, large black points with orange bars show the site-level response and 95% credible intervals (CrIs). The small points show population-specific responses; blue colored points denote credible population-level effects whereas grey colored points indicate cases where 95% CrIs for population effects include 0.

We found a consistent credible negative site-level effect of spring maximum temperature on TO at all sites: CP (−1.26, 95% CrI: -1.52 to - 1.01), DP (−1.73, 95% CrI: -1.99 to -1.49), LC (−1.23, 95% CrI: -1.48 to -1), SV (−0.66, 95% CrI: -0.9 to -0.43), and WA (−1.2, 95% CrI: -1.48 to -0.94). Negative species-specific effects were also observed across all species with credible species-specific effects within each site. This suggests that higher spring maximum temperatures are associated with earlier flight seasons across all sites and for many species (Figure 4 d). Winter precipitation had a credible positive site-level effect on TO at higher elevation sites, CP (0.71, 95% CrI: 0.48 to 0.94) and DP (0.85, 95% CrI: 0.64 to 1.06). This positive effect was also consistent across all species with credible species-specific effects at both sites suggesting that increased winter precipitation is associated with later flight periods at these two high-elevation sites. We found heterogeneity in the effect of winter precipitation within species at other sites (Figure 4 e). The effect of spring minimum temperature on timing was also heterogeneous within sites, except at LC, where we found a credible negative site-level effect, indicating that higher spring minimum temperatures are associated with earlier flight periods (Figure 4 f).

In general, we did not find credible site-level or consistent species-specific effects of climatic variables on LO (Figure 4 g, h, i). Nonetheless, a modest subset of species exhibited credible effects of each climate variable on LO at each of the five sites. In addition, at LC we found a credible positive site-level effect of spring maximum temperature (0.43, 95% CrI: 0.20 to 0.67), suggesting that higher spring maximum temperatures were associated with a longer season for most species (Figure 4 g). Conversely, at WA, we found a credible negative site-level effect of spring minimum temperature (−0.39, 95% CrI: -0.63 to -0.14), indicating that higher spring minimum temperatures were associated with shorter seasons (Figure 4 i).

### 3.3 Heterogeneous effects of climate on flight period

We analyzed variation in the effect of climate on different aspects of the occurrence distribution (MO, TO, and LO) at three levels: variation in the average effect of a climatic variable across sites, variation in the species-specific effects among species within a site averaged across sites, and variation in species-specific effects for each species among sites (populations) averaged across species. The results revealed substantial heterogeneity in the effects of all climatic variables (spring maximum temperature, winter precipitation, and spring minimum temperature) on each facet of the probability of occurrence distribution (Figure 5). The greatest heterogeneity was found among species within sites, followed by populations of the same species across different sites, with the least heterogeneity observed for mean-effects across sites (Figure 5).

**FIGURE 5.**
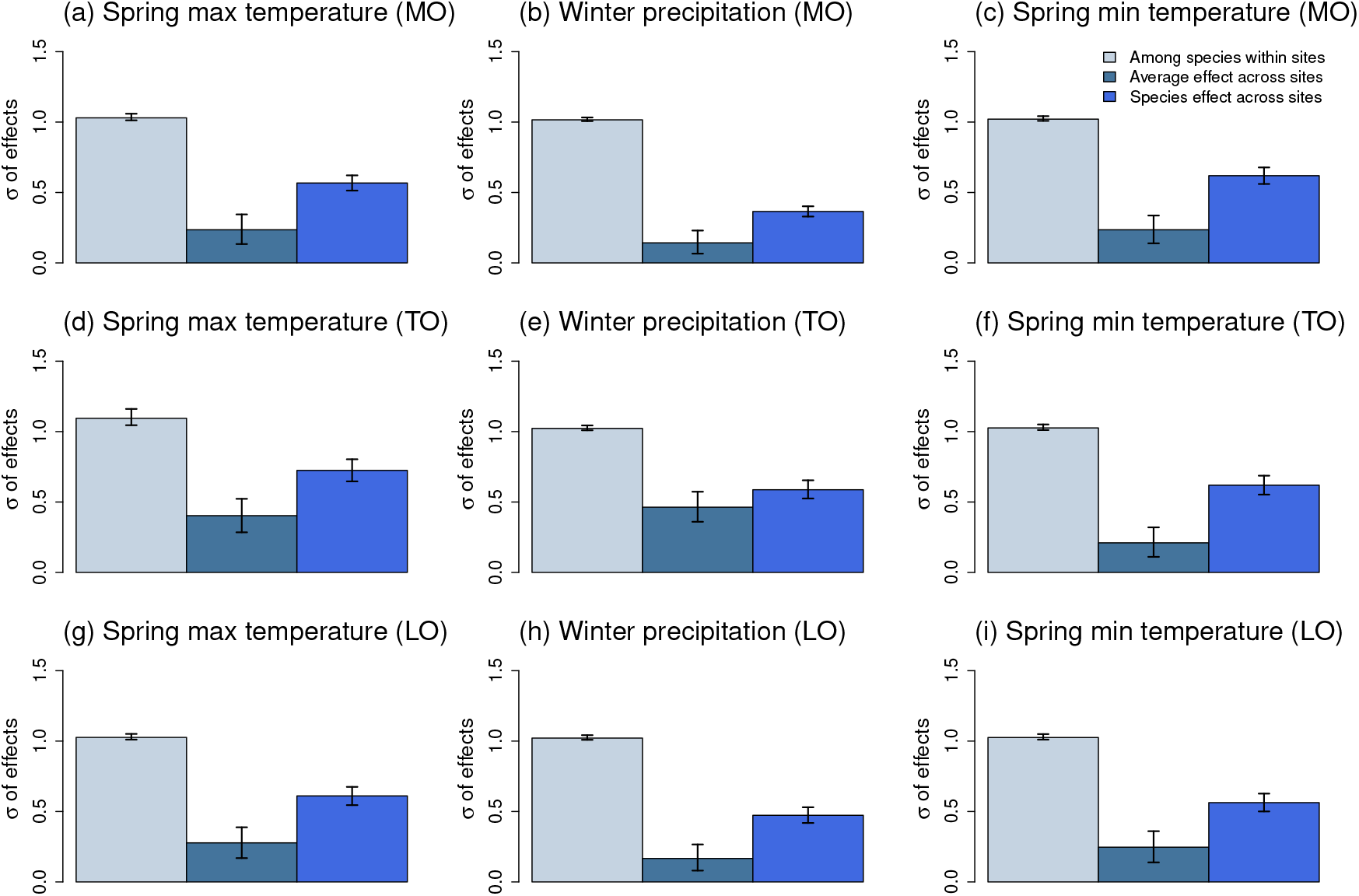
Variability in the effect of each climatic variable on butterfly flight period among species within a site (among species within site), across all sites (average effect across sites), and among populations of a species across sites (species effect across site). Results are shown for the effect of each climatic variable on the (a) mid-season occurrence (MO), (b) timing of occurrence (TO) and (c) length of occurrence (LO). Colored bars denote the estimated standard deviations (*σ* of effects) and error bars denote 95% confidence intervals on these estimates. Higher standard deviation indicates higher variability. The greatest heterogeneity was found among species within sites, followed by populations of the same species across different sites, with the least heterogeneity observed for mean-effects across sites.

The site-level heterogeneity in climate effects that were detected were not solely due to differences in species composition across sites (Figure A36). Specifically, with the exception of CP versus DP, which responded similarly to climate overall (Figure 4), species-level effects of climate on flight period were only weakly correlated for pairs of sites (Figure A36 a, b, d, e). In other words, the same climatic factor often had distinct effects on a given species at different sties.

### 3.4 Relationship between the effects of climate on different aspects of the occurrence distribution

To determine the extent to which the species-specific effect of each climatic variable on one facet of the occurrence distribution was associated with its effect on others, we estimated correlations between the species-specific effects of each climatic variable on MO and TO, MO and LO, and TO and LO at each site. Overall, we found that correlations across sites were weak such that climatic factors have largely independent effects on different aspects of the flight period (Figure 6). The main exceptions to the general weak correlations were as follows. We found moderate positive correlations between the effect of spring maximum temperature on MO and TO at CP (0.35, 95% CI: 0.12 to 0.55) and DP (0.32, 95% CI: 0.12 to 0.50), suggesting that the effects of spring maximum temperature on MO and TO covary more strongly at high elevations. Similarly, we detected moderate correlations in the effect of spring maximum temperature on TO and LO at CP (0.37, 95% CI: 0.13 to 0.56) and DP (0.56, 95% CI: 0.40 to 0.69), again indicating higher covariance at higher elevations. Lastly, there were moderate negative correlations in the effect of winter precipitation on MO and LO at CP, DP, LC, and SV (CP: -0.35, 95% CI: -0.55 to -0.11, DP: -0.35, 95% CI: -0.52 to -0.16, LC: -0.34, 95% CI: -0.52 to -0.14, and SV: -0.32, 95% CI: -0.51 to -0.09) (Figure 6).

**FIGURE 6.**
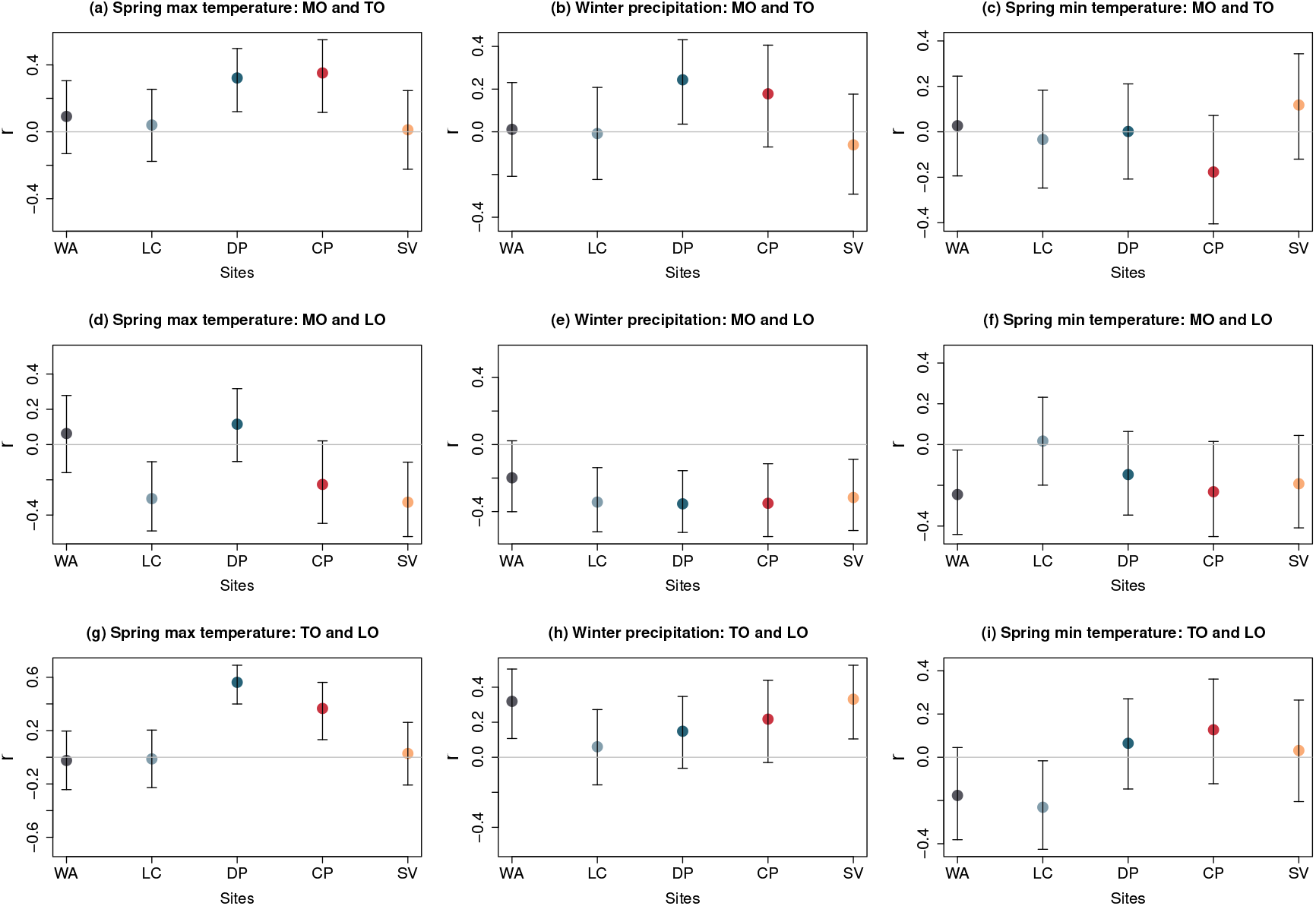
Relationship between the effect of each climatic variable on different aspects of the occurrence distribution of species at each site as captured by the Pearson correlation coefficient (*r*). The panels show (a) the correlation between the effects of spring maximum temperature on the mid-season occurrence and timing of occurrence (MO and TO), (b) the correlation between the effects of winter precipitation on the mid-season occurrence and timing of occurrence (MO and TO), (c) the correlation between the effects of spring minimum temperature on the mid-season occurrence and timing of occurrence (MO and TO), (d) the correlation between the effects of spring maximum temperature on the mid-season occurrence and length of occurrence (MO and LO), (e) the correlation between the effects of winter precipitation on the mid-season occurrence and length of occurrence (MO and LO), (f) the correlation between the effects of spring minimum temperature on the mid-season occurrence and length of occurrence (MO and LO), (g) the correlation between the effects of spring maximum temperature on the timing and length of occurrence (TO and LO), (h) the correlation between the effects of winter precipitation on the timing and length of occurrence (TO and LO), and (i) the correlation between the effects of spring minimum temperature on the timing and length of occurrence (TO and LO). The correlation coefficient indicates the similarity in the effect of each climatic variable on two compared aspects of the occurrence distribution of species at each site.

### 3.5 Multivariate analysis of climate effects on flight period

To simultaneously assess the influence of multiple climatic variables and examine how their species-specific effects on different facets of the occurrence distribution are correlated, we used a PCA. This approach enabled us to move beyond pairwise correlations and detect broader patterns of how each climate variable impacts various aspects of the occurrence distribution simultaneously. At CP and DP (the highest elevation sites), the first principal component explained variation in the effect of climate on TO and MO, while the second principal component (PCA 2) captured variation in climatic effects on LO (Figure 7 a, b). This indicates that the effects of climate on flight season length diverge from its effects on the timing and abundance of butterflies at these sites. Interestingly, this pattern was not detected at the lower elevation sites–LC, SV, and WA (Figure 7 c, d, e), suggesting site-specific associations.

**FIGURE 7.**
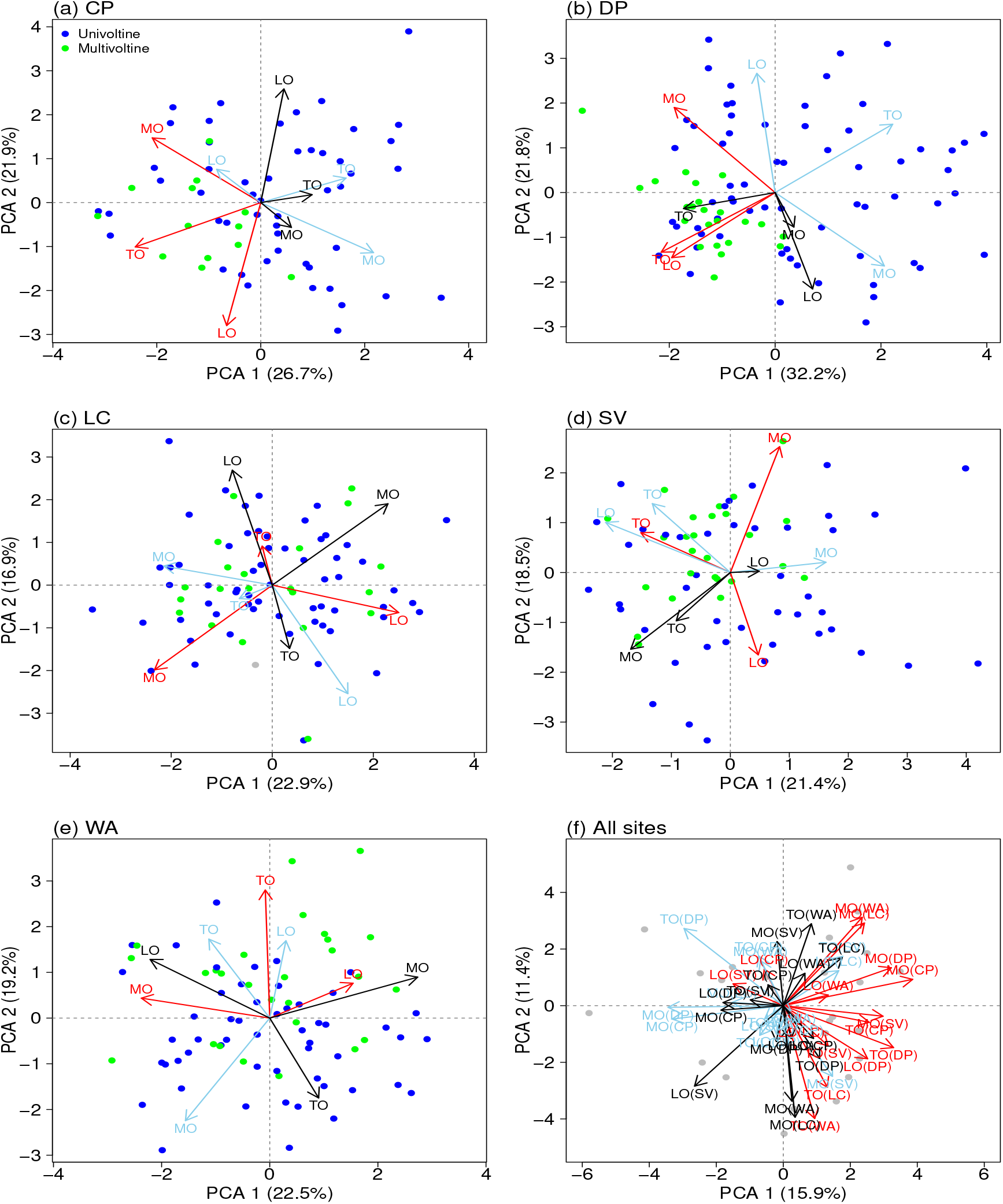
Principal component analysis (PCA) of the effects of all climate variables on different aspects of the flight period (mid-season occurrence (MO), timing of occurrence (TO), length of occurrence (LO)) across species observed at (a) Castle Peak, (b) Donner Pass, (c) Lang Crossing, Sierra Valley, (e) Washington, and (f) all sites. Red arrows represent the effect of spring maximum temperature, blue arrows represent the effect of winter precipitation, and black arrows represent the effect of spring minimum temperature. Blue points represent univoltine species while green points represent multivoltine species.

We found that at CP, DP, SV, and WA, the relationship between the effect of climate on different aspects of the flight period differed between multivoltine and univoltine species, as they formed distinct clusters in the PCA plots (Figure 7a, b, d, e) with Euclidean distances of 1.56, 1.87, 0.97 and 1.22 at CP, DP, SV and WA respectively. A permutation test confirmed that these differences were statistically significant and not due to chance (p < 0.05; Figure A37a, b, d, e). This suggests that voltinism may influence, or otherwise be associated with how effects of climate are realized on different aspect of the flight periods. However, this pattern was not observed at LC (Figure 7c, A37c).

Finally we examine whether the associations between the effects of climate on different aspects of the flight period show similar patterns across sites. Our analysis showed that patterns of association were often site-specific. However, these patterns were more similar between geographically proximate sites, such as CP and DP, and LC and WA (Figure 7f).

## 4 DISCUSSION

Climate change is having widespread consequences for the phenology of wild plants and animals (Cohen et al. 2018, Forister et al. 2018, Newson et al. 2016, Parmesan 2006 2007). Long-term datasets that span diverse habitats and elevation gradients, such as the dataset analyzed in this study, provide critical insights into the effects of climate on phenology. By examining both site- and species-level responses, we quantified the distinct effects of climate at both the species and community levels. Consistent with prior studies, we found that the flight periods of butterfly species are changing over time (Colom et al. 2021, Habel et al. 2024). Furthermore, our results indicate declines in MO over time across all sites, suggesting that butterfly populations are becoming increasingly scarce and less abundant. Similar results have been documented in previous studies of data from our focal sites that modeled abundance without considering phenology (that is, studies that did not allow the probability of occurrence to vary across the season within years) (Halsch et al. 2024).

Our findings complement and extend earlier research reinforcing the idea that climate is a major driver of phenological shifts in butterfly populations. Specifically, our models identified spring maximum temperature, winter precipitation, and spring minimum temperature as the most influential climatic predictors (among those considered) of butterfly flight period across all sites. TO was particularly sensitive to spring maximum temperatures, with warmer temperatures resulting in earlier emergence at all sites. This finding is consistent with previous studies that report earlier emergence of butterfly species under warming conditions (Colom et al. 2021, Forister and Shapiro 2003, Gutiérrez and Wilson 2021, Habel et al. 2024, Parmesan 2007), likely due to increased developmental rates at higher temperatures. At higher elevation sites– CP and DP–we observed a pronounced effect of winter precipitation (which usually comes in the form of snow) on TO, with increased precipitation leading to delayed TO. Slower snowmelt at these elevations could act as a thermal buffer, insulating the ground and maintaining cooler temperatures that slow the development of butterflies and their resources, such as host plants and nectar sources (Inouye 2008). This finding highlights the critical role of winter precipitation, particularly in montane habitats, emphasizing the idea that shifts in precipitation patterns are as crucial as temperature changes in shaping ecological processes. This is consistent with previous studies that have demonstrated strong responses of insect populations to changes in snow cover (Boggs and Inouye 2012, Halsch et al. 2024).

In addition, our results underscore the substantial influence of even modest increases in nighttime (daily minimum) temperatures on the flight period. In some cases, studies on insect phenology or abundance have focused on daily maximum temperature or average daily temperatures, often overlooking the impact of nighttime temperatures, which are increasing more rapidly due to climate change at these sites and in many other regions (Vasseur et al. 2014, Speights et al. 2017). Our findings suggest that nighttime warming may elicit different phenological responses compared to daytime warming, potentially affecting not only developmental rates but also ecological interactions. Moreover, species exhibit varying responses to extreme daily conditions, which are often obscured when using average daily temperatures as a metric (Ma et al. 2015). By considering both minimum and maximum daily temperatures, rather than relying solely on averages, we can obtain a more nuanced understanding of ecological responses to unusual weather events, daily fluctuations, or specific temperature thresholds, which are often critical but underappreciated in phenology studies.

We observed the highest heterogeneity in the effect of climate among species within sites, followed by populations of the same species across different sites, with the least heterogeneity observed for the average effect of climate across sites. This suggests that, while climate change is a broad driver of shifts in phenology, species-specific traits (such as life history and behavioral traits) modulate differences in climate responses (Colom et al. 2021, Diamond et al. 2011, Faltýnek Fric et al. 2020, Gutiérrez and Wilson 2021). The substantial variation among populations of the same species across different sites could result from differences in the range of climatic conditions experienced at each location or may indicate that these populations are locally adapted to their respective environments.

Our analysis further reveals that climate affects different facets of the occurrence distribution in distinct ways at each site. The observed differences in site-specific patterns were not solely due to variations in species composition between sites, suggesting that local environmental conditions and microclimatic factors may play a critical role in shaping how climate drives changes in butterfly occurrence distributions at different locations. Importantly, MO, TO, and LO did not respond uniformly to climate variables, reinforcing the limitations of focusing exclusively on metrics such as the day of first flight. For instance, while TO may advance with warming temperatures, LO and MO might remain stable or even increase, complicating predictions of population dynamics and species interactions. Such complexities can confound meta-analyses that rely on different metrics to interpret species-level phenological shifts. Additionally, they can lead to over-interpretation or obscuring of the full extent of phenological changes, which may have important implications for understanding mismatches in species interactions driven by climate change. Additionally, we find that voltinism could influences how climate effects on different facets of the flight period are related. Our findings complement and extend earlier research suggesting that the life history traits of species play a significant role in determining their phenological responses to climate (Diamond et al. 2011, Zografou et al. 2021). Future studies focusing on how various natural history traits, such as voltinism, diapause strategies, and diet breadth, modulate the effects of climate on butterfly phenology would further enhance our understanding of both species-specific and community-level responses to climate change.

Our findings emphasize the importance of considering multiple facets of phenology and climate drivers when assessing the impacts of climate change on species phenology. Conservation strategies must recognize that populations of the same species may respond differently to climate change depending on their location and ecological context. We acknowledge that some species observed at these sites are migratory and do not directly experience winter or spring weather conditions at these sites, as they overwinter in different environments. However, climate patterns are increasingly becoming more spatially and temporally auto-correlated in the face of climate change, making it plausible for climate conditions in the study area to have indirect associations with offsite climate effects (Di Cecco and Gouhier 2018, Liebhold et al. 2004) (also see Figure 2 c, d, e). Finally, this study highlights the complexity of predicting climate-driven changes in phenology and stresses the need for species- and site-specific approaches, as local adaptation could buffer or exacerbate the effects of global climate trends on a regional scale. By adopting a more holistic view of phenological responses, we can gain deeper insights into how species respond to changing climatic conditions, better inform conservation strategies, and improve our ability to predict future phenological patterns in a warming world.

## AUTHOR CONTRIBUTIONS

**Gbolahan A Reis**: Conceptualization (equal); Data Curation (equal); Formal Analysis (lead); Funding Acquisition (equal); Methodology (equal); Visualization (lead); Writing – Original Draft Preparation (lead); Writing – Review & Editing (equal). **Matthew L Forister**: Conceptualization(equal); Data Curation (equal); Investigation (equal); Funding Acquisition (equal); Writing – Review & Editing (equal). **Christopher A Halsch**: Data Curation (equal); Investigation (equal); Writing – Review & Editing (equal). **Clare M Dittemore**: Data Curation (equal); Investigation (equal); Writing – Review & Editing (equal). **Arthur M Shapiro**: Data Curation (equal); Investigation (equal); Writing – Review & Editing (equal). **Zachariah Gompert**: Conceptualization (equal); Data Curation (equal); Funding Acquisition (equal); Methodology (equal); Supervision (lead); Writing – Review & Editing (equal).

## ACKNOWLEDGMENTS

We gratefully acknowledge the support and resources provided by the Center for High Performance Computing at the University of Utah. This work was also supported by the National Science Foundation (DEB-2114974 to Z.G., DEB-2114973 to M.L.F.), the Utah State University Ecology Center Research Award (A07339-1092), and the American Philosophical Society grant to G.A.R.

## APPENDIX A SUPPLEMENTARY MATERIAL

