## Supplemental Table 1 and Figure 1-37 for "Temporal occupancy distributions reveal multifaceted and heterogeneous effects of climatic variation on montane butterflies"

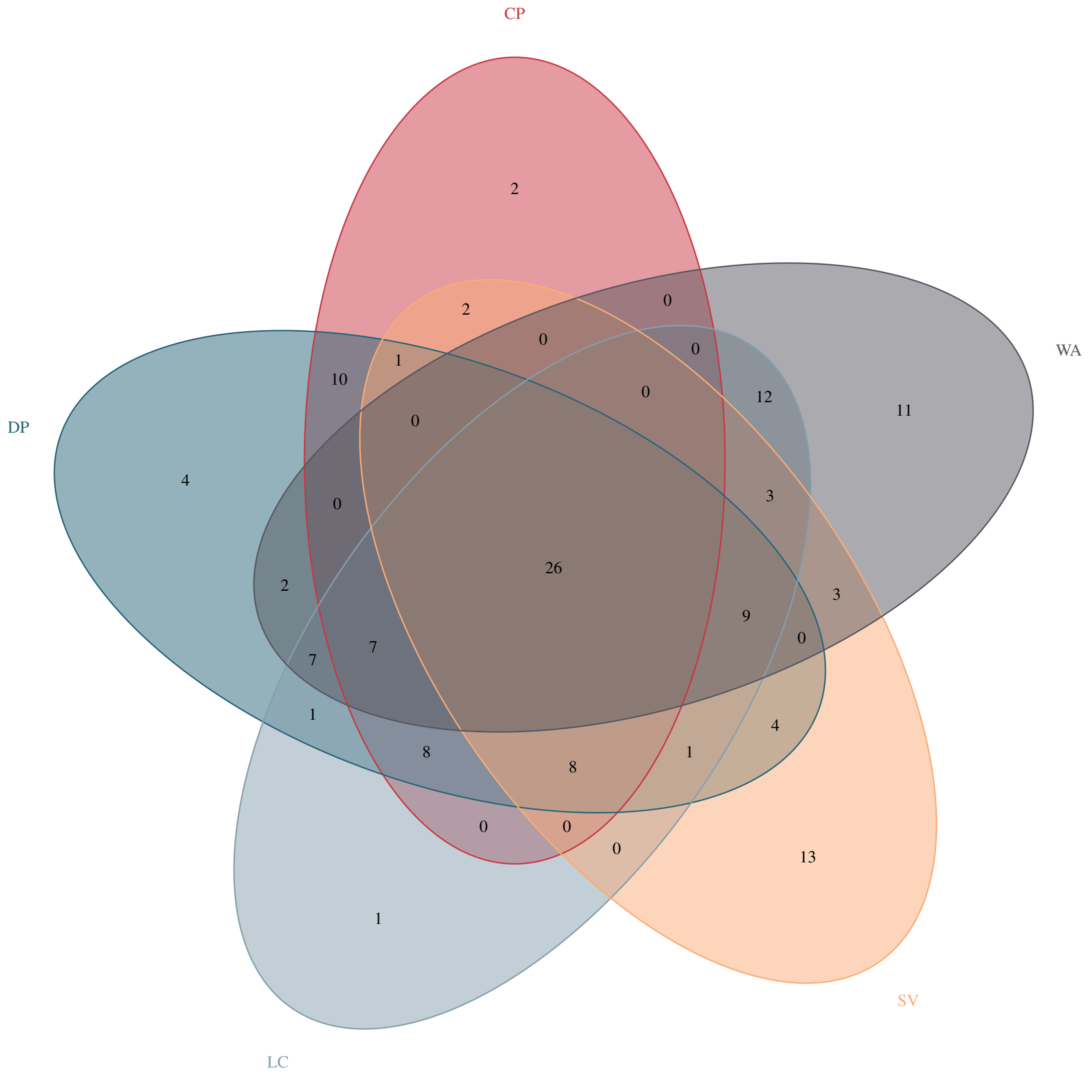


**FIGURE A1** Distribution of butterfly species across five monitoring sites. The Venn diagram illustrates the number of species shared between sites and the number of species common across all sites.

**TABLE** A1: Comparison of models using the forward selection approach. The expected log pointwise predictive density (ELPD) was estimated via leave-one-out cross-validation. ELPD values were averaged across all sites for each model, and the model with the highest average ELPD was identified as the best-performing model. Climate covariates are described as follows: OD refers to the ordinal day of observation in the current year; Year represents the current year; GDD corresponds to growing degree days accumulated from January 1 to May 31 of the current year. SWE indicates the mean daily snow water equivalent from September 1 of the previous year to May 31 of the current year. Fall variables include FMINT, FPREC, and FMAXT, which represent the mean daily minimum temperature, mean daily precipitation, and mean daily maximum temperature, respectively, from September 1 to November 30 of the previous year. Winter variables include WMINT, WPREC, and WMAXT, representing the mean daily minimum temperature, mean daily precipitation, and mean daily maximum temperature, respectively, from December 1 of the previous year to February 28 of the current year. Spring variables include SMINT, SPREC, and SMAXT, which correspond to the mean daily minimum temperature, mean daily precipitation, and mean daily maximum temperature, respectively, from March 1 to May 31 of the current year.

| Covariates | No | CP | DP | LC | SV | WA | Average |
|---|---|---|---|---|---|---|---|
| OD, Year | 0 | -8235.4 | -16317 | -14693.3 | -13555 | -13713.5 | -13302.84 |
| Gdd, OD, Year | 1 | -7708.7 | -14802.7 | -13880.1 | -13378.6 | -13414.7 | -12636.96 |

Continued on next page

**A1 – continued from previous page**

| Covariates | No | CP | DP | LC | SV | WA | Average |
|---|---|---|---|---|---|---|---|
| SWE, OD, Year | 1 | -8250.5 | -16349.8 | -14726.8 | -13596.8 | -13747.6 | -13334.3 |
| FMINT, OD, Year | 1 | -8248.6 | -16213.7 | -14635.8 | -13576.5 | -13726.8 | -13280.28 |
| FPREC, OD, Year | 1 | -8196.8 | -16305.1 | -14769.4 | -13618.5 | -13796.8 | -13337.32 |
| FMAXT, OD, Year | 1 | -8216.8 | -16263.7 | -14716.1 | -13638.7 | -13742.7 | -13315.6 |
| WMINT, OD, Year | 1 | -8265.1 | -16325.1 | -14713.3 | -13565.2 | -13745.4 | -13322.82 |
| WPREC, OD, Year | 1 | -8053.8 | -15987.8 | -14692.9 | -13575.8 | -13767.4 | -13215.54 |
| WMAXT, OD, Year | 1 | -8172 | -16154.9 | -14679.2 | -13623.1 | -13719 | -13269.64 |
| SMINT, OD, Year | 1 | -8000.2 | -15486 | -14078.8 | -13476 | -13454.2 | -12899.04 |
| SPREC, OD, Year | 1 | -7873.3 | -15358 | -14172.8 | -13461.1 | -13545.1 | -12882.06 |
| SMAXT, OD, Year | 1 | -7609.8 | -14742.4 | -13830 | -13323.2 | -13326.5 | -12566.38 |
| SMAXT, Gdd, OD, Year | 2 | -7656.5 | -14712.1 | -13789.1 | -13355.5 | -13362.5 | -12575.14 |
| SMAXT, SWE, OD, Year | 2 | -7679.1 | -14752.4 | -13871.2 | -13344.3 | -13336.9 | -12596.78 |
| SMAXT, SPREC, OD, Year | 2 | -7659.2 | -14726.9 | -13875.3 | -13342.6 | -13385.4 | -12597.88 |
| SMAXT, SMINT, OD, Year | 2 | -7567.7 | -14703.4 | -13813.3 | -13317 | -13278.3 | -12535.94 |
| SMAXT, FMAXT, OD, Year | 2 | -7652.6 | -14733.2 | -13869.9 | -13397 | -13402 | -12610.94 |
| SMAXT, FPREC, OD, Year | 2 | -7619.5 | -14773.3 | -13922.4 | -13368.2 | -13389.3 | -12614.54 |
| SMAXT, FMINT, OD, Year | 2 | -7668.6 | -14727.6 | -13839.5 | -13331.7 | -13359.8 | -12585.44 |
| SMAXT, WMAXT, OD, Year | 2 | -7594 | -14693.6 | -13888 | -13388.9 | -13363.2 | -12585.54 |
| SMAXT, WPREC, OD, Year | 2 | -7379.6 | -14475.5 | -13856.4 | -13324.9 | -13360.8 | -12479.44 |
| SMAXT, WMINT, OD, Year | 2 | -7673 | -14749.1 | -13863.1 | -13320.7 | -13329.9 | -12587.16 |
| SMAXT, WPREC, FMAXT, OD, Year | 3 | -7470.5 | -14503.4 | -13908.3 | -13406.8 | -13437.4 | -12545.28 |
| SMAXT, WPREC, FPREC, OD, Year | 3 | -7438.1 | -14535.8 | -13955.5 | -13375.4 | -13429.1 | -12546.78 |
| SMAXT, WPREC, FMINT, OD, Year | 3 | -7441.7 | -14420 | -13872.8 | -13336.4 | -13393.8 | -12492.94 |
| SMAXT, WPREC, Gdd, OD, Year | 3 | -7447 | -14478.6 | -13815.7 | -13345.2 | -13410.4 | -12499.38 |
| SMAXT, WPREC, SWE, OD, Year | 3 | -7465.5 | -14454 | -13901.9 | -13347 | -13377.5 | -12509.18 |
| SMAXT, WPREC, SPREC, OD, Year | 3 | -7440.4 | -14470.1 | -13902.3 | -13342.2 | -13420.8 | -12515.16 |
| SMAXT, WPREC, SMINT, OD, Year | 3 | -7377.1 | -14402.7 | -13834.1 | -13321.6 | -13318.1 | -12450.72 |
| SMAXT, WPREC, WMAXT, OD, Year | 3 | -7367.3 | -14320.9 | -13892.2 | -13357.9 | -13333.6 | -12462.26 |
| SMAXT, WPREC, WPREC, OD, Year | 3 | -7381.6 | -14441.2 | -13903.5 | -13310.1 | -13321.2 | -12441.52 |
| SMAXT, WPREC, FMAXT, FPREC, OD, Year | 4 | -7360.1 | -14419.5 | -13899.8 | -13343.1 | -13330.4 | -12454.62 |
| SMAXT, WPREC, FMINT, FPREC, OD, Year | 4 | -7374.5 | -14400.9 | -13884.8 | -13337.8 | -13333.6 | -12442.24 |
| SMAXT, WPREC, FMAXT, SWE, OD, Year | 4 | -7406.9 | -14446.9 | -13909.2 | -13336.4 | -13347.3 | -12485.68 |
| SMAXT, WPREC, FMINT, SWE, OD, Year | 4 | -7354.2 | -14408.1 | -13895.4 | -13335.9 | -13354.6 | -12445.24 |
| SMAXT, WPREC, WMINT, SWE, OD, Year | 4 | -7353.9 | -14388.5 | -13892.7 | -13328.9 | -13349.3 | -12442.46 |
| SMAXT, WPREC, WMAXT, SWE, OD, Year | 4 | -7314.3 | -14358.5 | -13834.2 | -13336.4 | -13336.1 | -12426.76 |

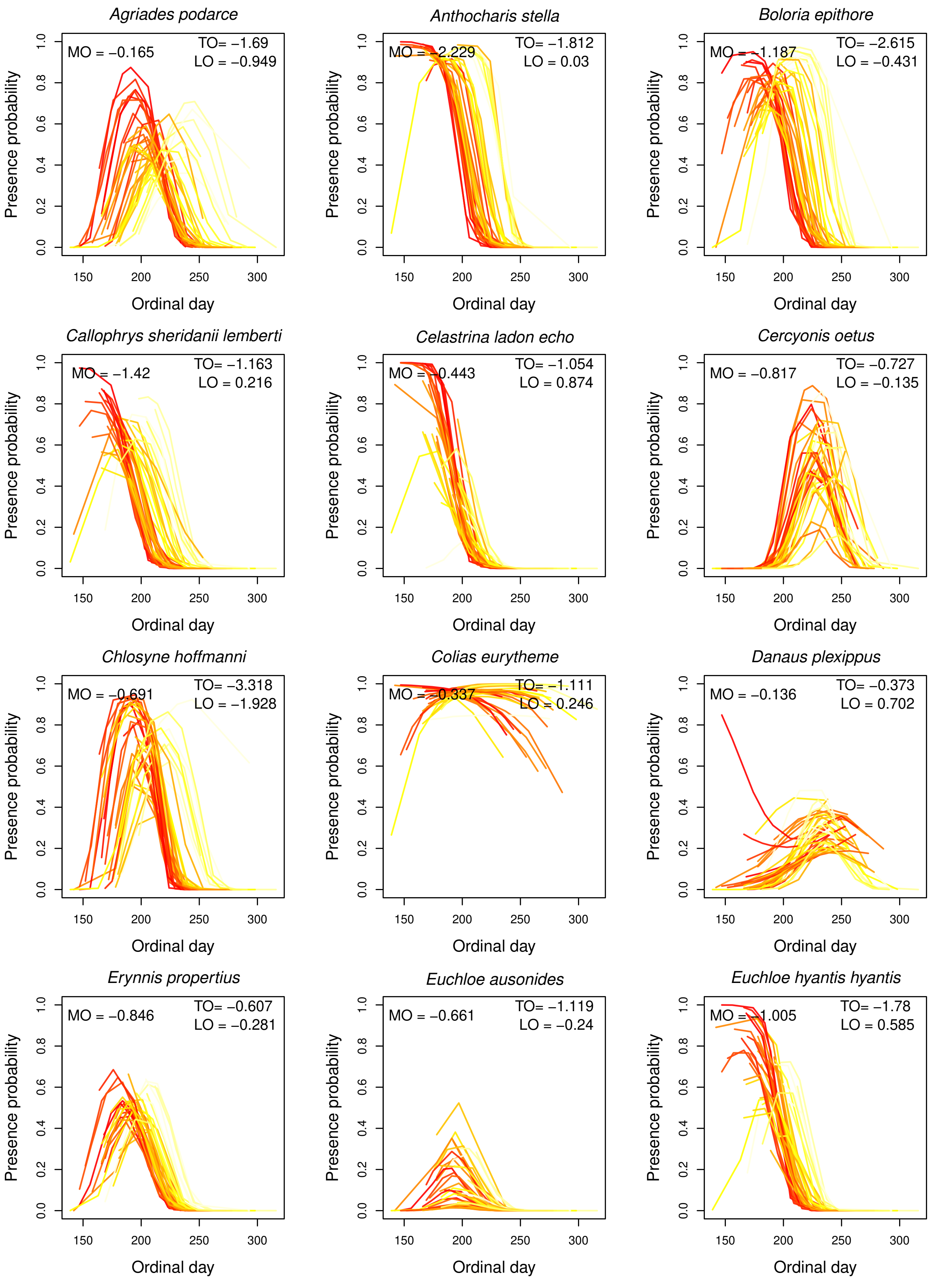


**FIGURE A2** Estimates of interannual variation in flight periods of species at Castle Peak from our climate model. Each line represents the probability of occurrence on each day across a year, with colors indicating average spring maximum temperatures for that year (darker colors indicate higher spring maximum temperatures). Thus the effects of spring maximum temperature on MO, TO and LO for each species are shown [illegible]

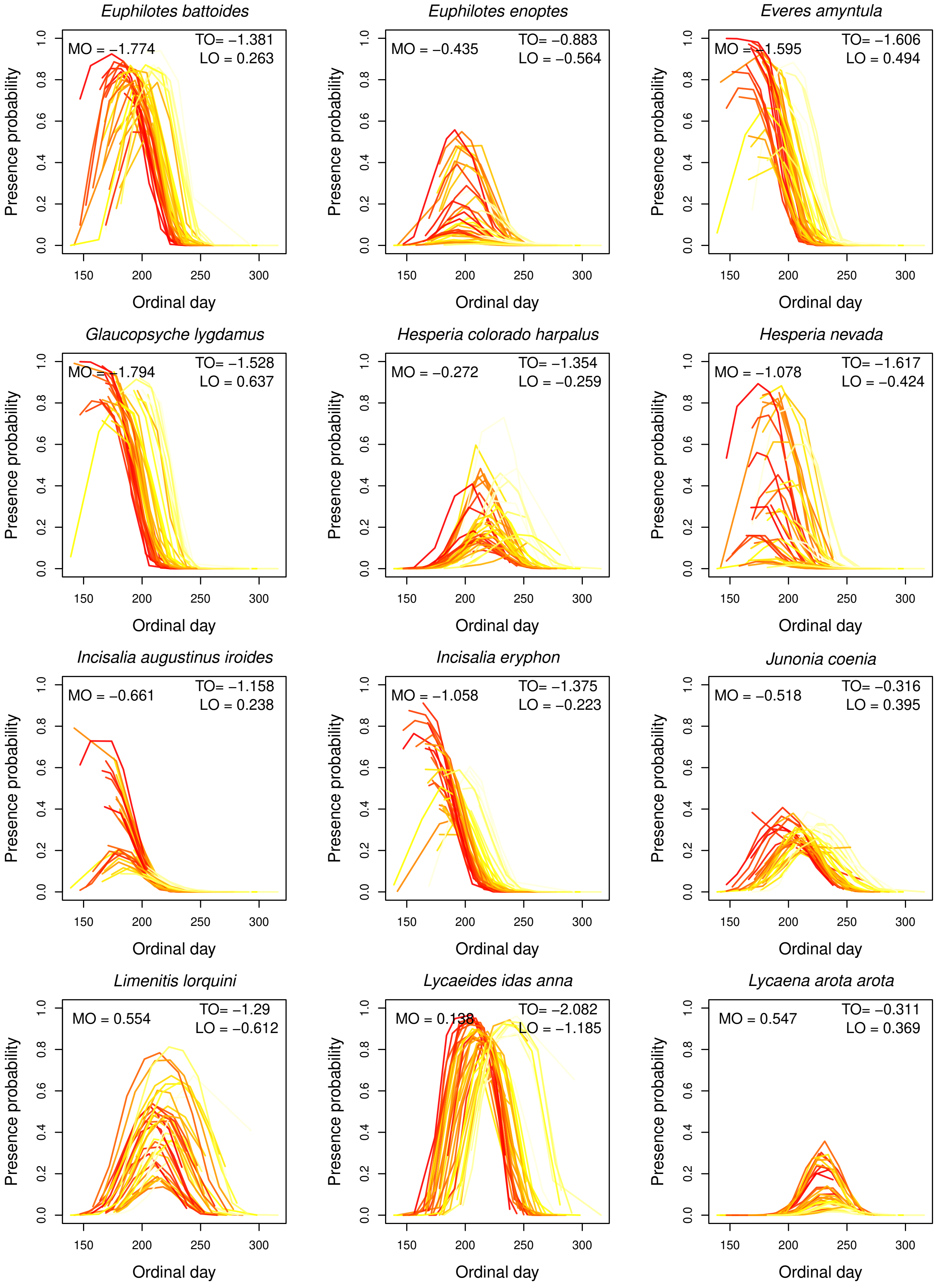


**FIGURE A3** Estimates of interannual variation in flight periods of species at Castle Peak from our climate model. Each line represents the probability of occurrence on each day across a year, with colors indicating average spring maximum temperatures for that year (darker colors indicate higher spring maximum temperatures). Thus the effects of spring maximum temperature on MO, TO and LO for each species are shown [illegible]

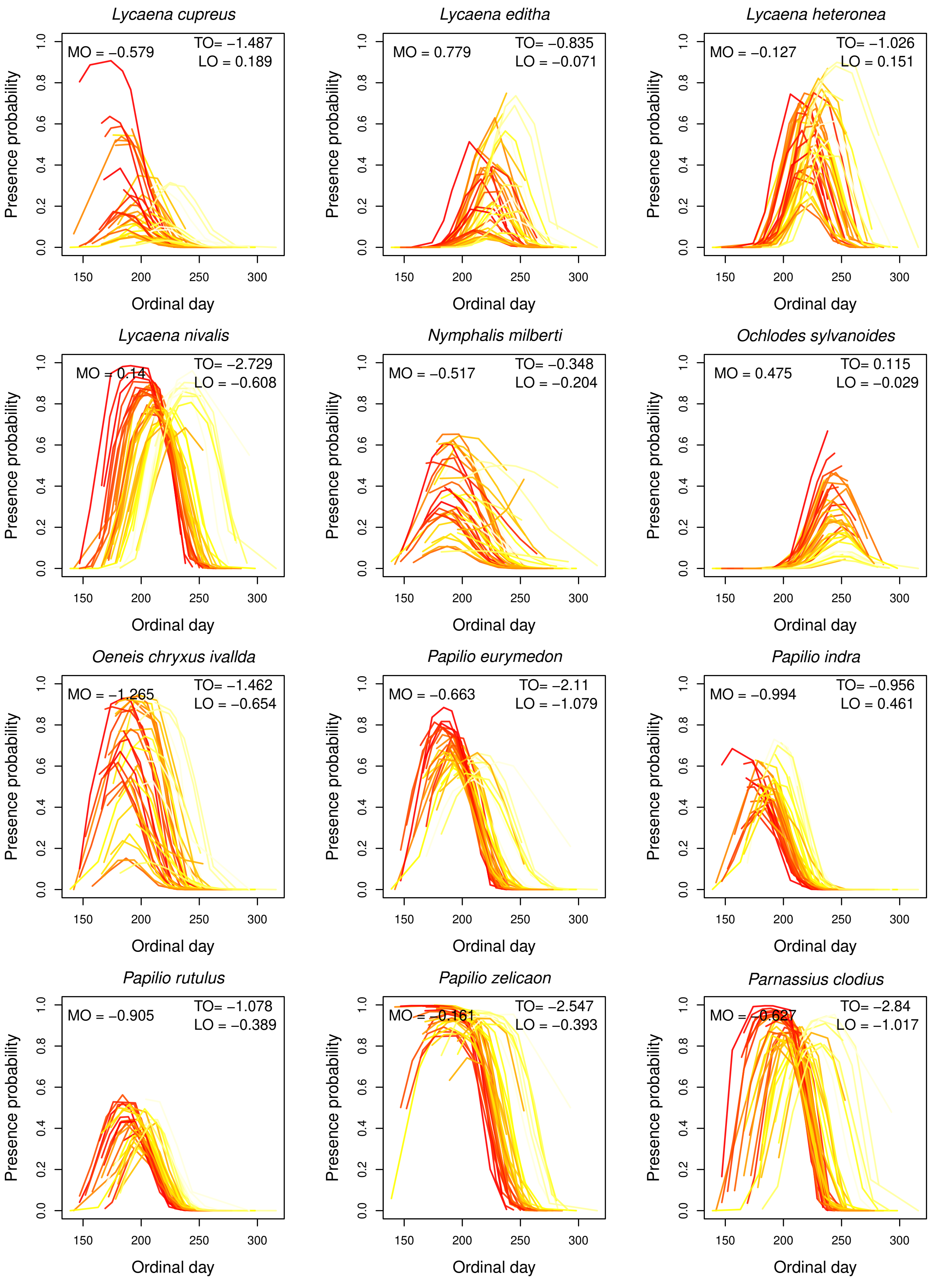


**FIGURE A4** Estimates of interannual variation in flight periods of species at Castle Peak from our climate model. Each line represents the probability of occurrence on each day across a year, with colors indicating average spring maximum temperatures for that year (darker colors indicate higher spring maximum temperatures). Thus the effects of spring maximum temperature on MO, TO and LO for each species are shown [illegible]

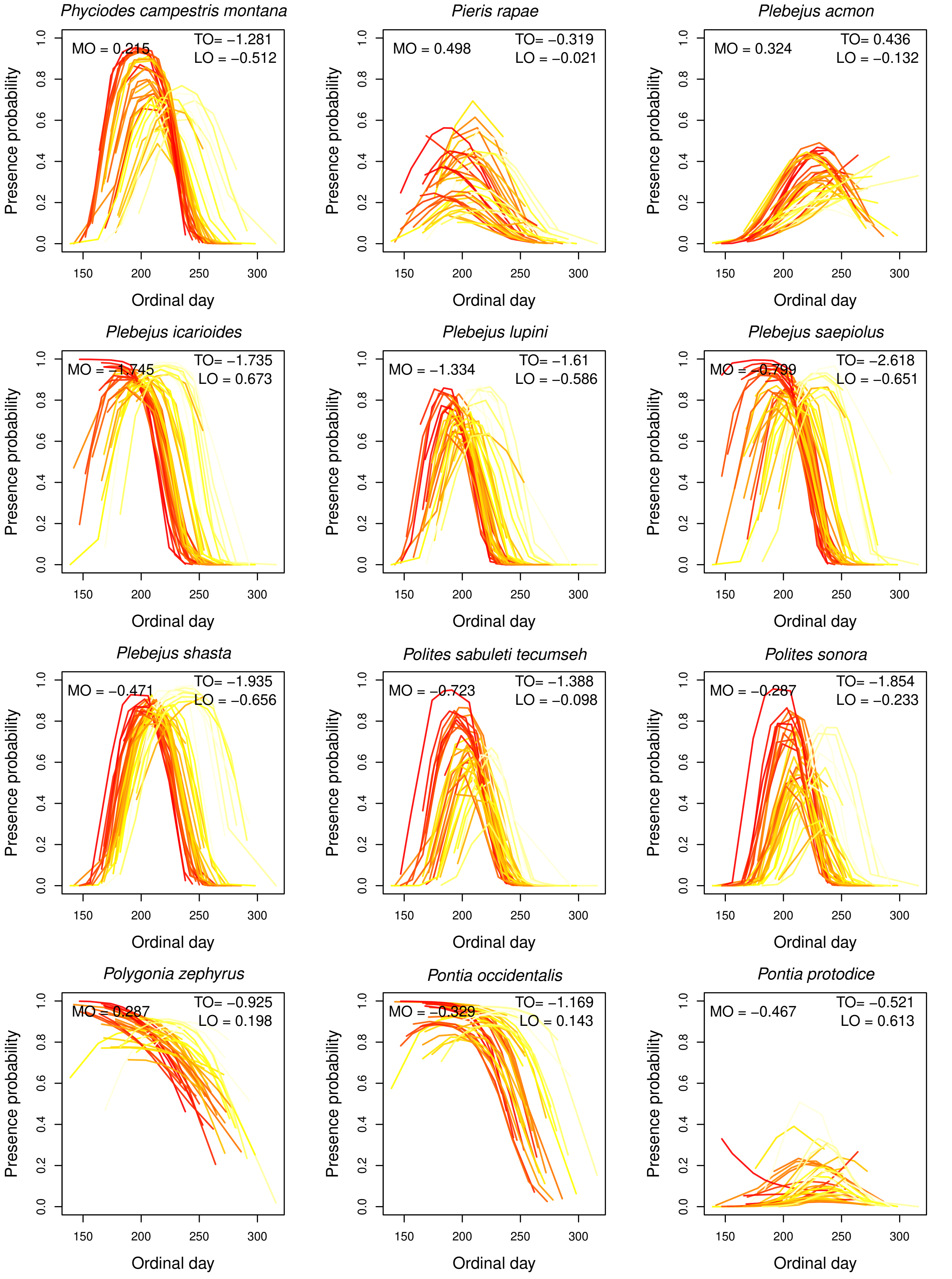


**FIGURE A5** Estimates of interannual variation in flight periods of species at Castle Peak from our climate model. Each line represents the probability of occurrence on each day across a year, with colors indicating average spring maximum temperatures for that year (darker colors indicate higher spring maximum temperatures). Thus the effects of spring maximum temperature on MO, TO and LO for each species are shown [illegible]

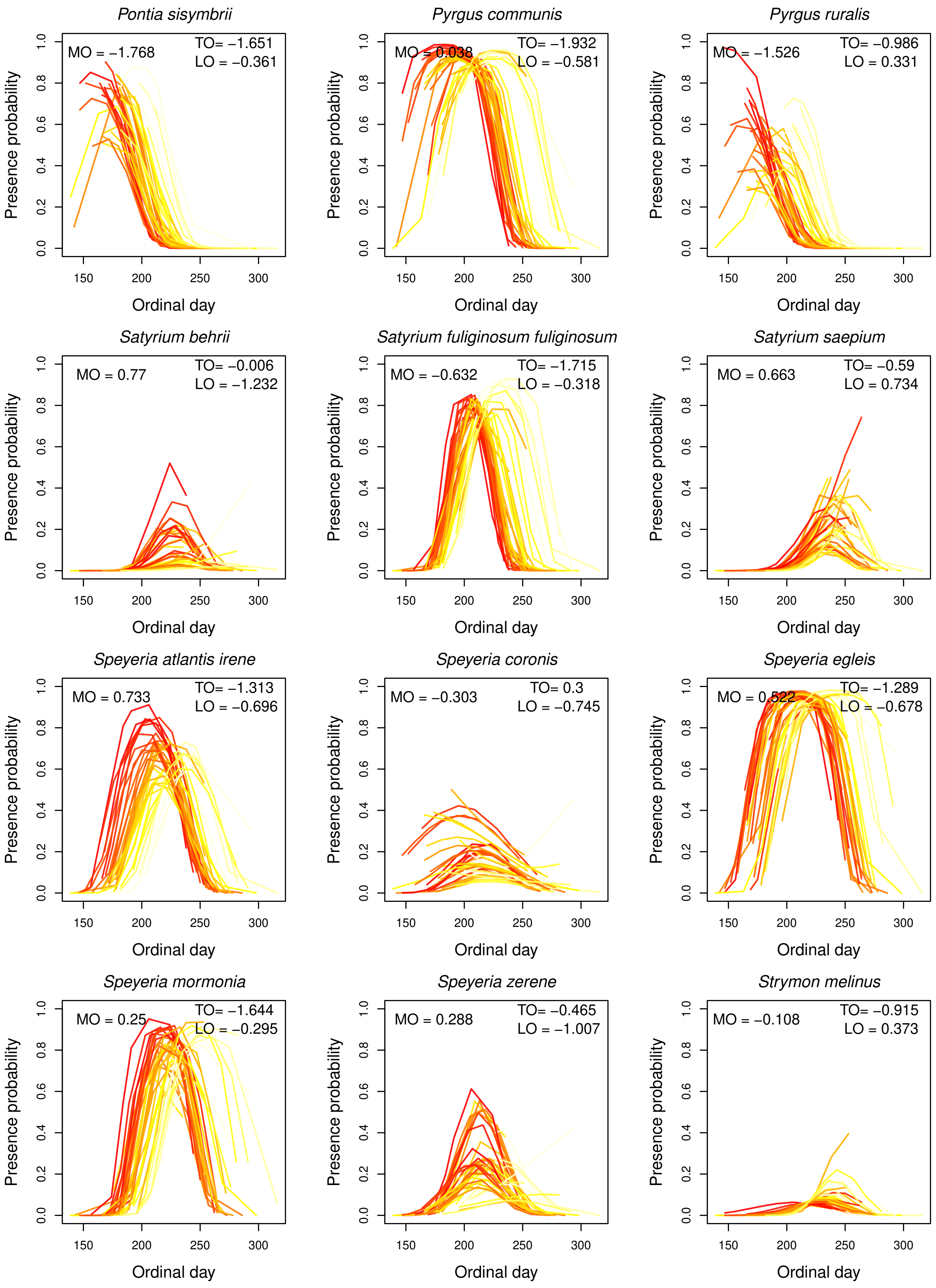


**FIGURE A6** Estimates of interannual variation in flight periods of species at Castle Peak from our climate model. Each line represents the probability of occurrence on each day across a year, with colors indicating average spring maximum temperatures for that year (darker colors indicate higher spring maximum temperatures). Thus the effects of spring maximum temperature on MO, TO and LO for each species are shown [illegible]

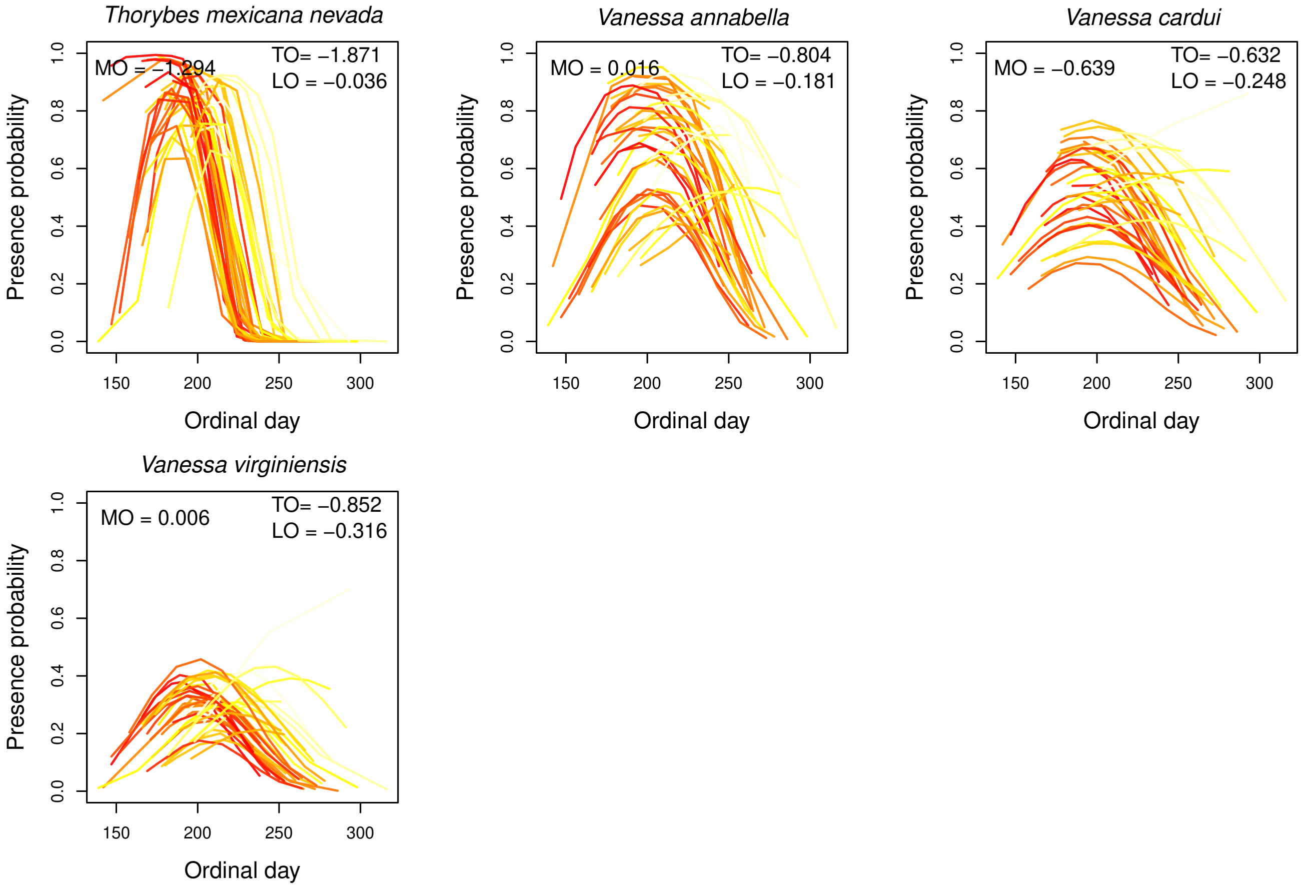


**FIGURE A7** Estimates of interannual variation in flight periods of species at Castle Peak from our climate model. Each line represents the probability of occurrence on each day across a year, with colors indicating average spring maximum temperatures for that year (darker colors indicate higher spring maximum temperatures). Thus the effects of spring maximum temperature on MO, TO and LO for each species are shown [illegible]

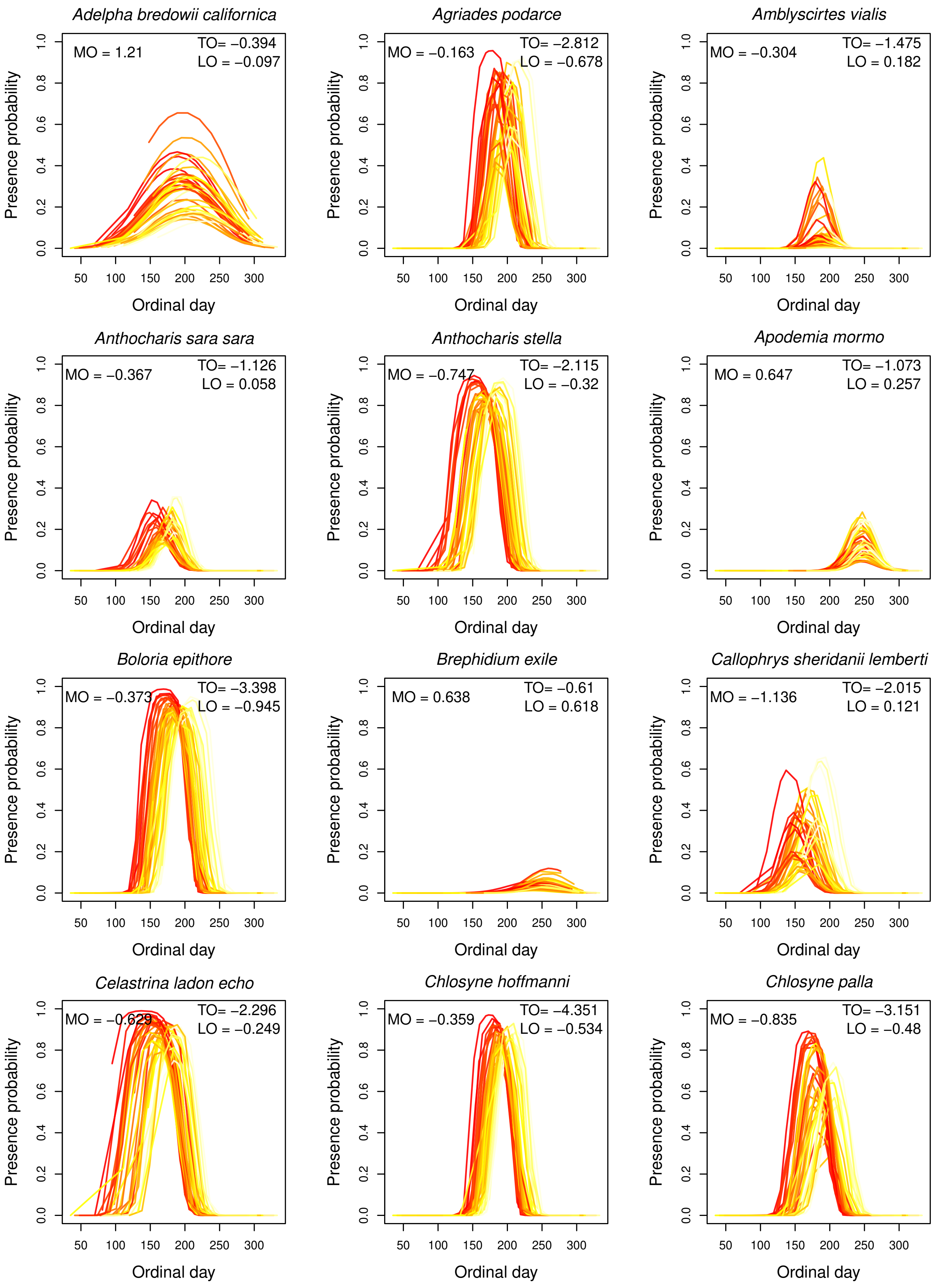


**FIGURE A8** Estimates of interannual variation in flight periods of species at Donner Pass from our climate model. Each line represents the probability of occurrence on each day across a year, with colors indicating average spring maximum temperatures for that year (darker colors indicate higher spring maximum temperatures). Thus the effects of spring maximum temperature on MO, TO and LO for each species are shown in each plot.

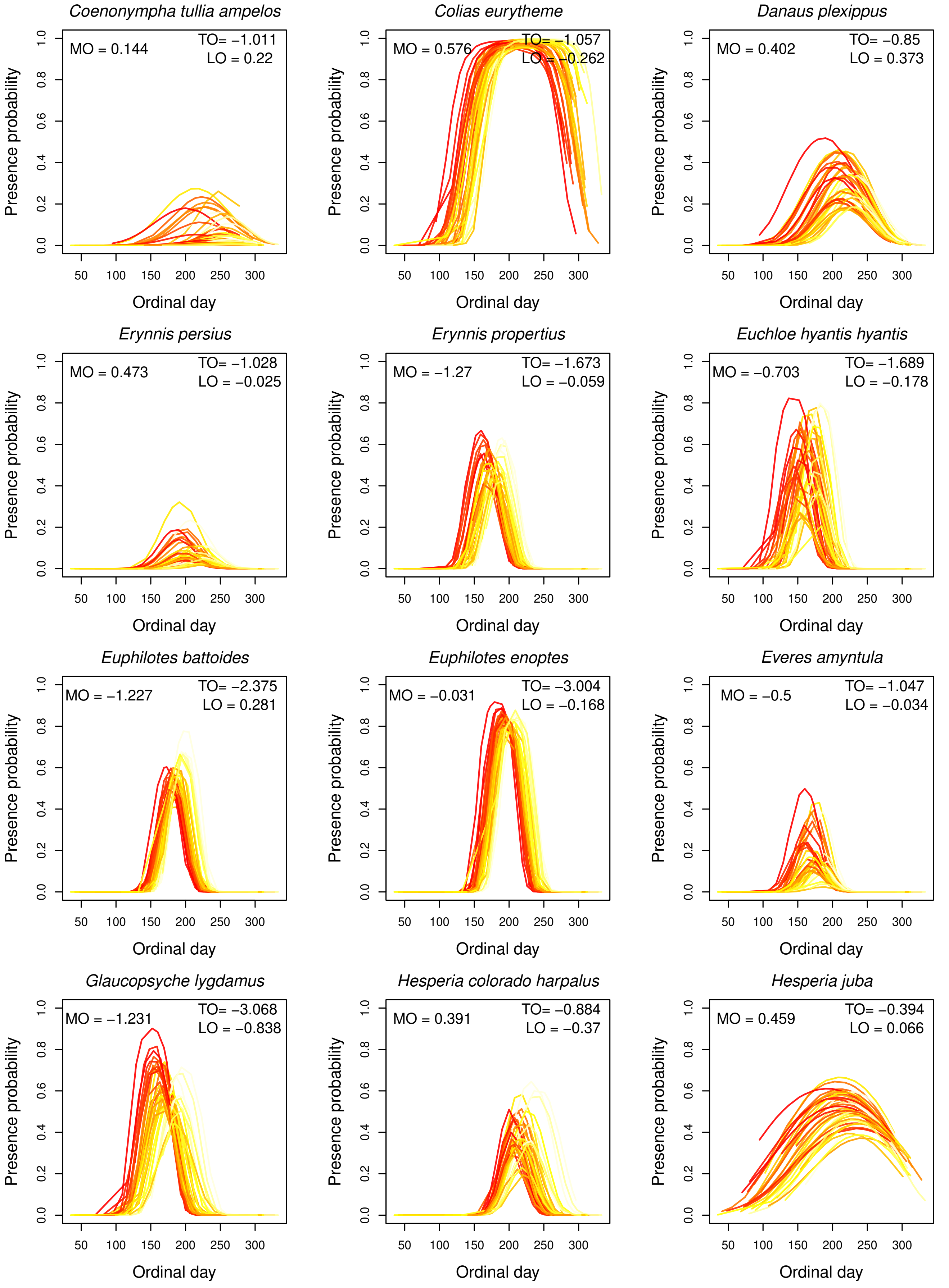


**FIGURE A9** Estimates of interannual variation in flight periods of species at Donner Pass from our climate model. Each line represents the probability of occurrence on each day across a year, with colors indicating average spring maximum temperatures for that year (darker colors indicate higher spring maximum temperatures). Thus the effects of spring maximum temperature on MO, TO and LO for each species are shown [illegible]

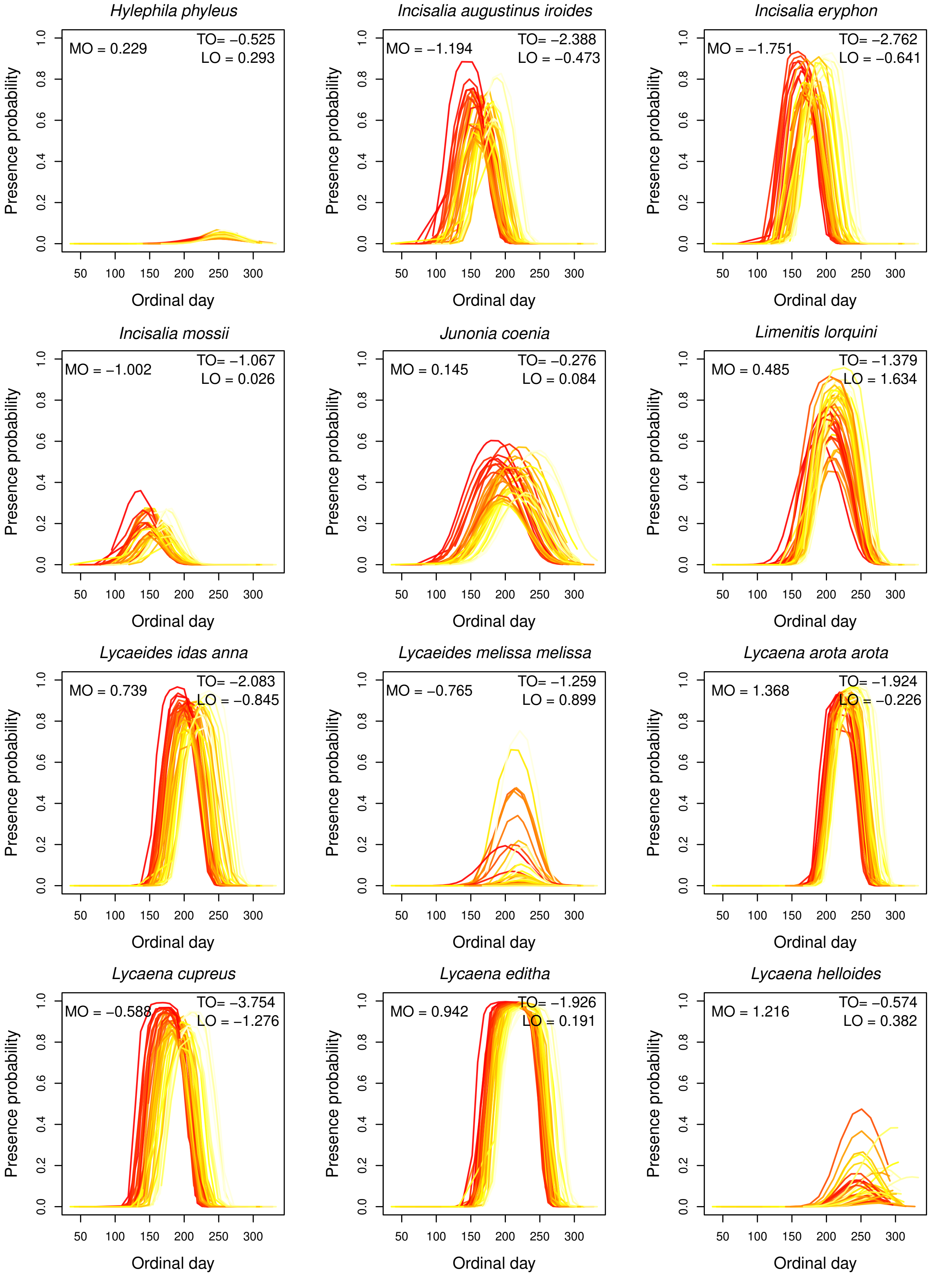


**FIGURE A10** Estimates of interannual variation in flight periods of species at Donner Pass from our climate model. Each line represents the probability of occurrence on each day across a year, with colors indicating average spring maximum temperatures for that year (darker colors indicate higher spring maximum temperatures). Thus the effects of spring maximum temperature on MO, TO and LO for each species are shown [illegible]

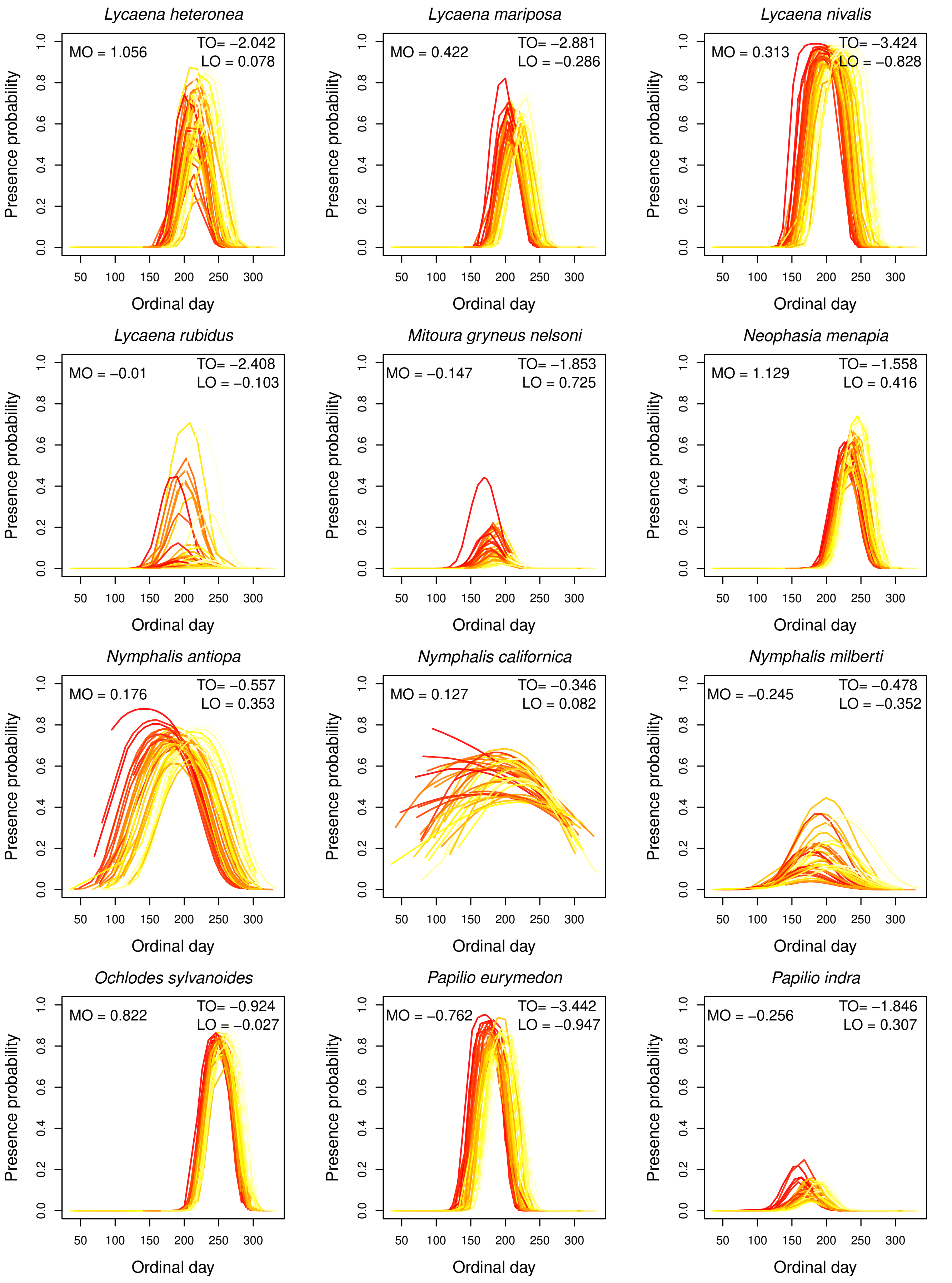


**FIGURE A11** Estimates of interannual variation in flight periods of species at Donner Pass from our climate model. Each line represents the probability of occurrence on each day across a year, with colors indicating average spring maximum temperatures for that year (darker colors indicate higher spring maximum temperatures). Thus the effects of spring maximum temperature on MO, TO and LO for each species are shown [illegible]

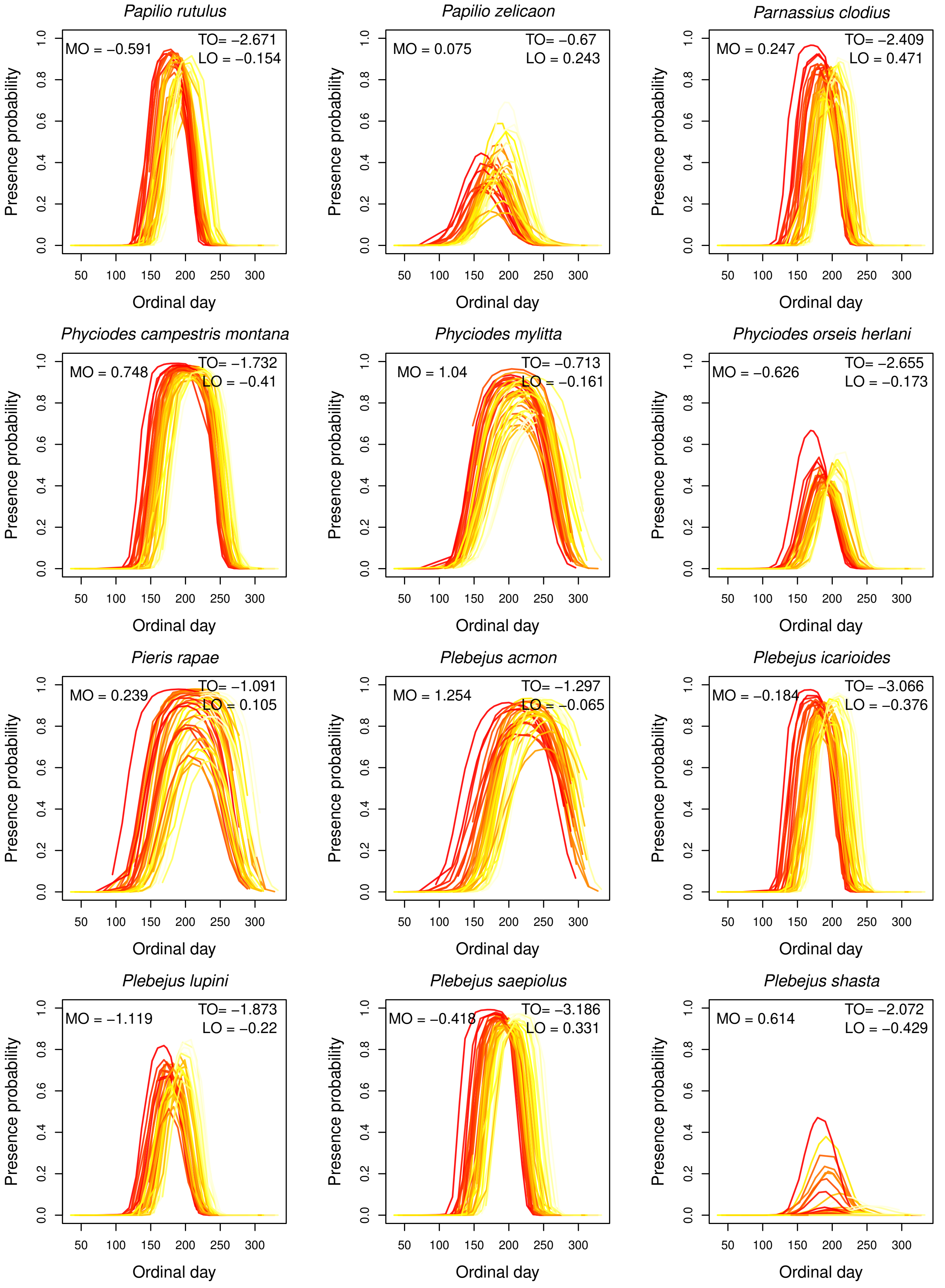


**FIGURE A12** Estimates of interannual variation in flight periods of species at Donner Pass from our climate model. Each line represents the probability of occurrence on each day across a year, with colors indicating average spring maximum temperatures for that year (darker colors indicate higher spring maximum temperatures). Thus the effects of spring maximum temperature on MO, TO and LO for each species are shown [illegible]

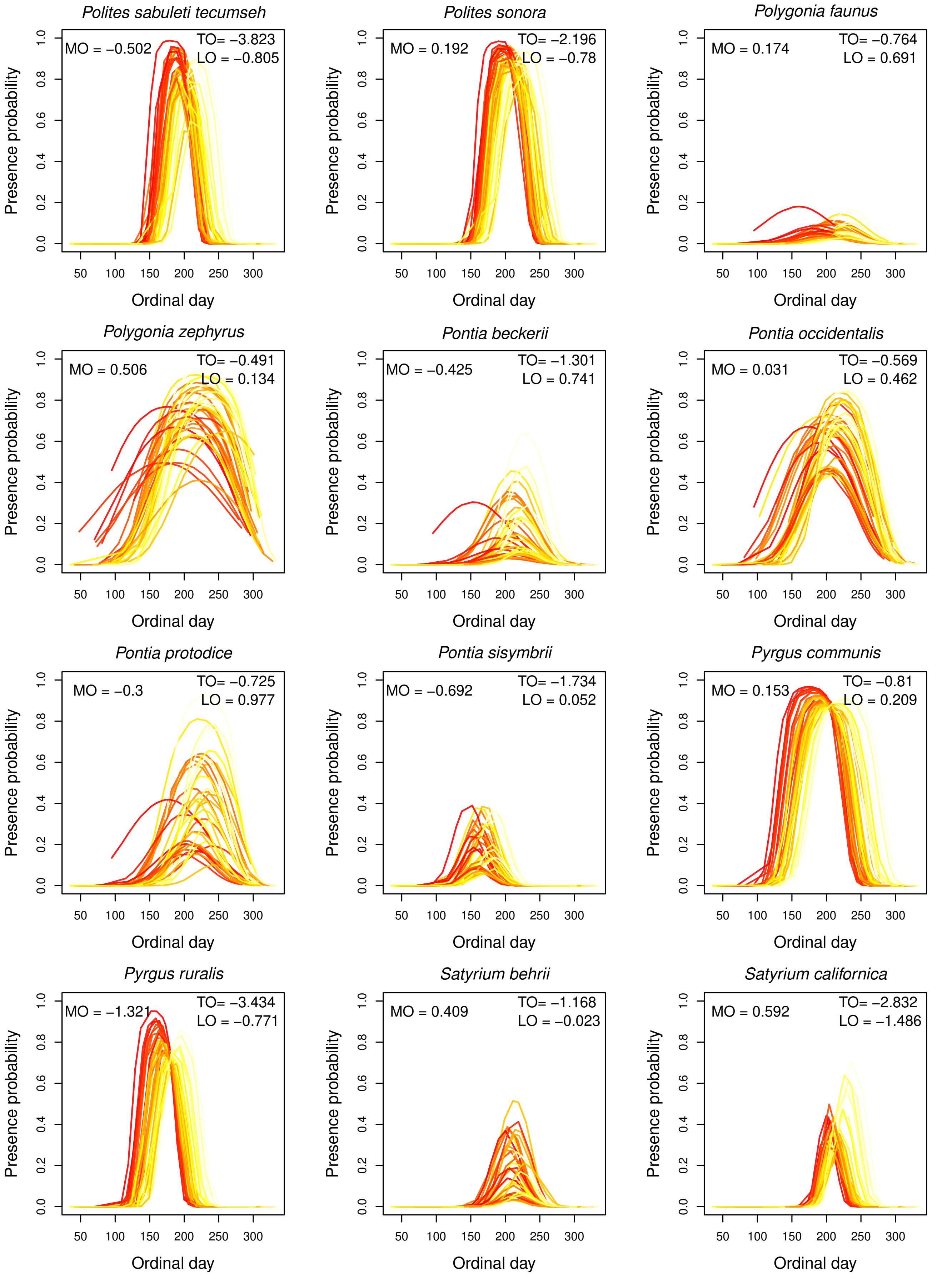


**FIGURE A13** Estimates of interannual variation in flight periods of species at Donner Pass from our climate model. Each line represents the probability of occurrence on each day across a year, with colors indicating average spring maximum temperatures for that year (darker colors indicate higher spring maximum temperatures). Thus the effects of spring maximum temperature on MO, TO and LO for each species are shown [illegible]

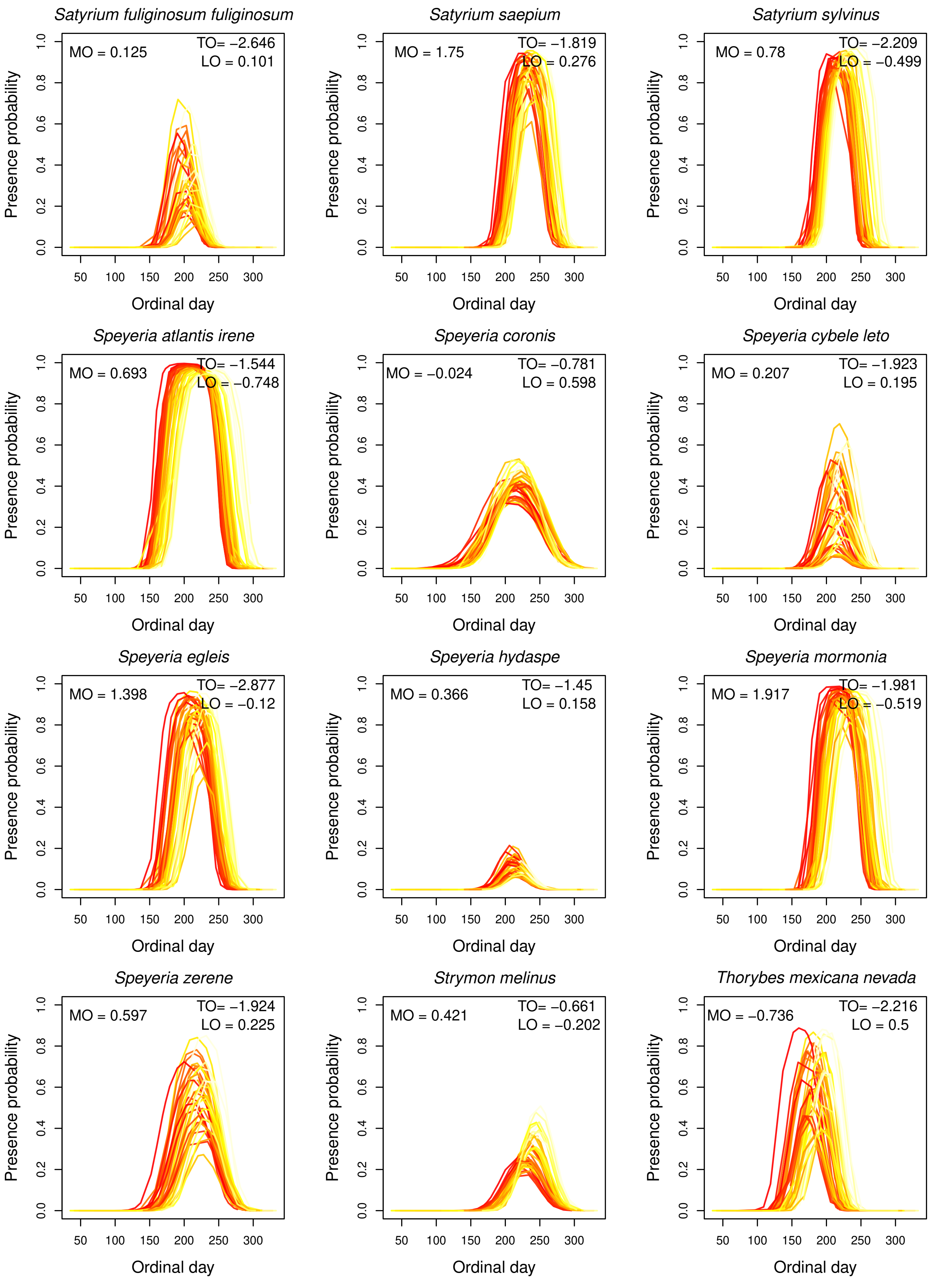


**FIGURE A14** Estimates of interannual variation in flight periods of species at Donner Pass from our climate model. Each line represents the probability of occurrence on each day across a year, with colors indicating average spring maximum temperatures for that year (darker colors indicate higher spring maximum temperatures). Thus the effects of spring maximum temperature on MO, TO and LO for each species are shown [illegible]

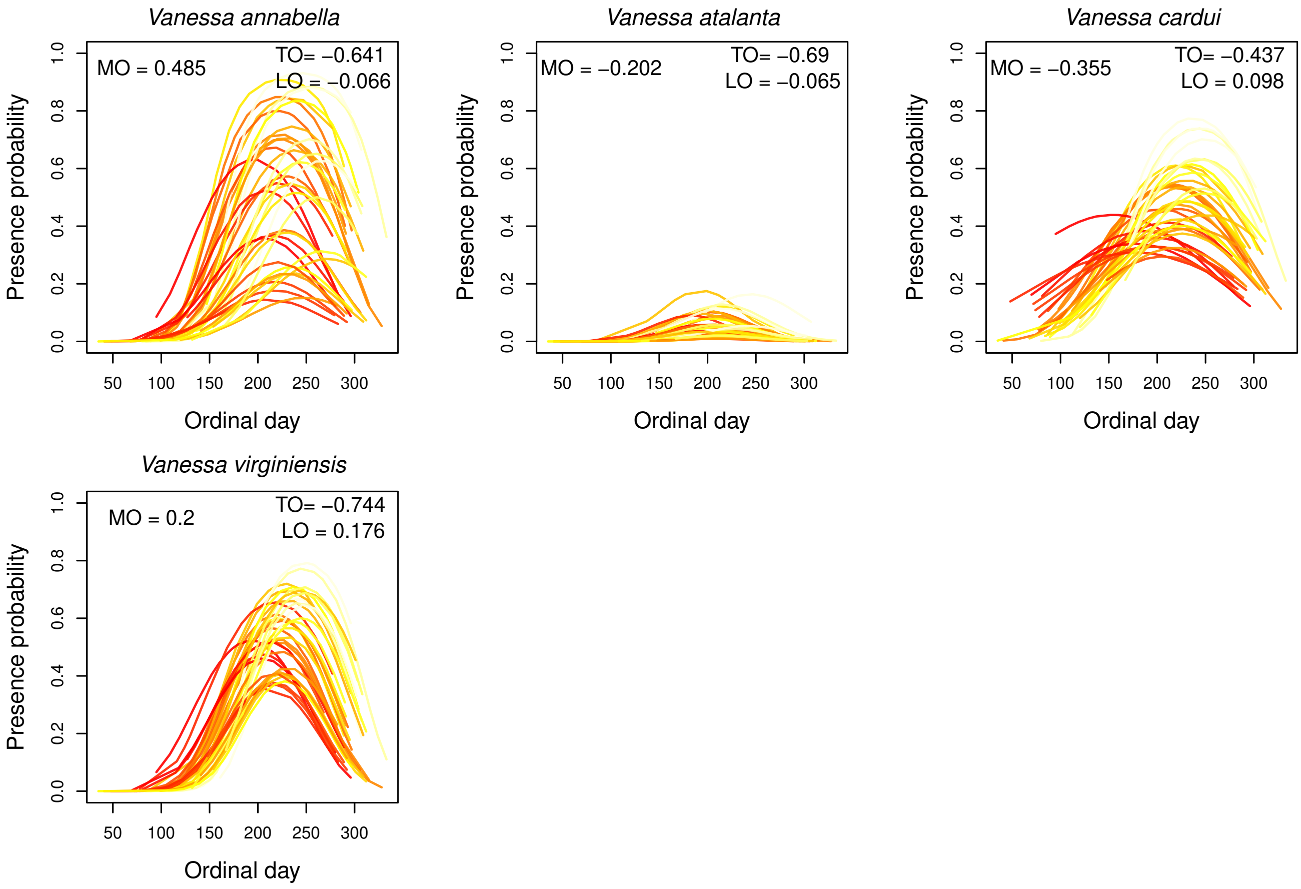


**FIGURE A15** Estimates of interannual variation in flight periods of species at Donner Pass from our climate model. Each line represents the probability of occurrence on each day across a year, with colors indicating average spring maximum temperatures for that year (darker colors indicate higher spring maximum temperatures). Thus the effects of spring maximum temperature on MO, TO and LO for each species are shown [illegible]

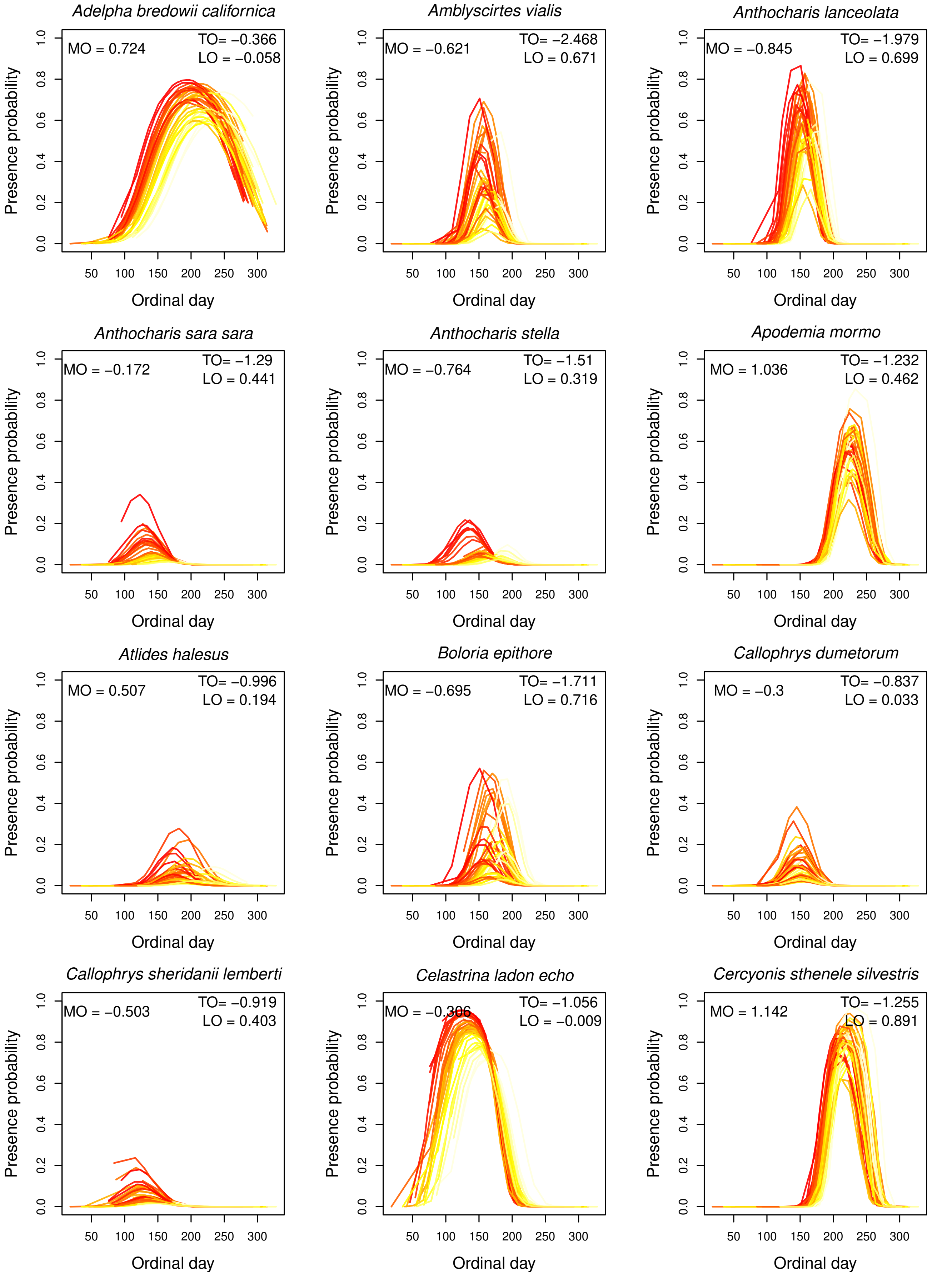


**FIGURE A16** Estimates of interannual variation in flight periods of species at Lang Crossing from our climate model. Each line represents the probability of occurrence on each day across a year, with colors indicating average spring maximum temperatures for that year (darker colors indicate higher spring maximum temperatures). Thus the effects of spring maximum temperature on MO, TO and LO for each species are shown [illegible]

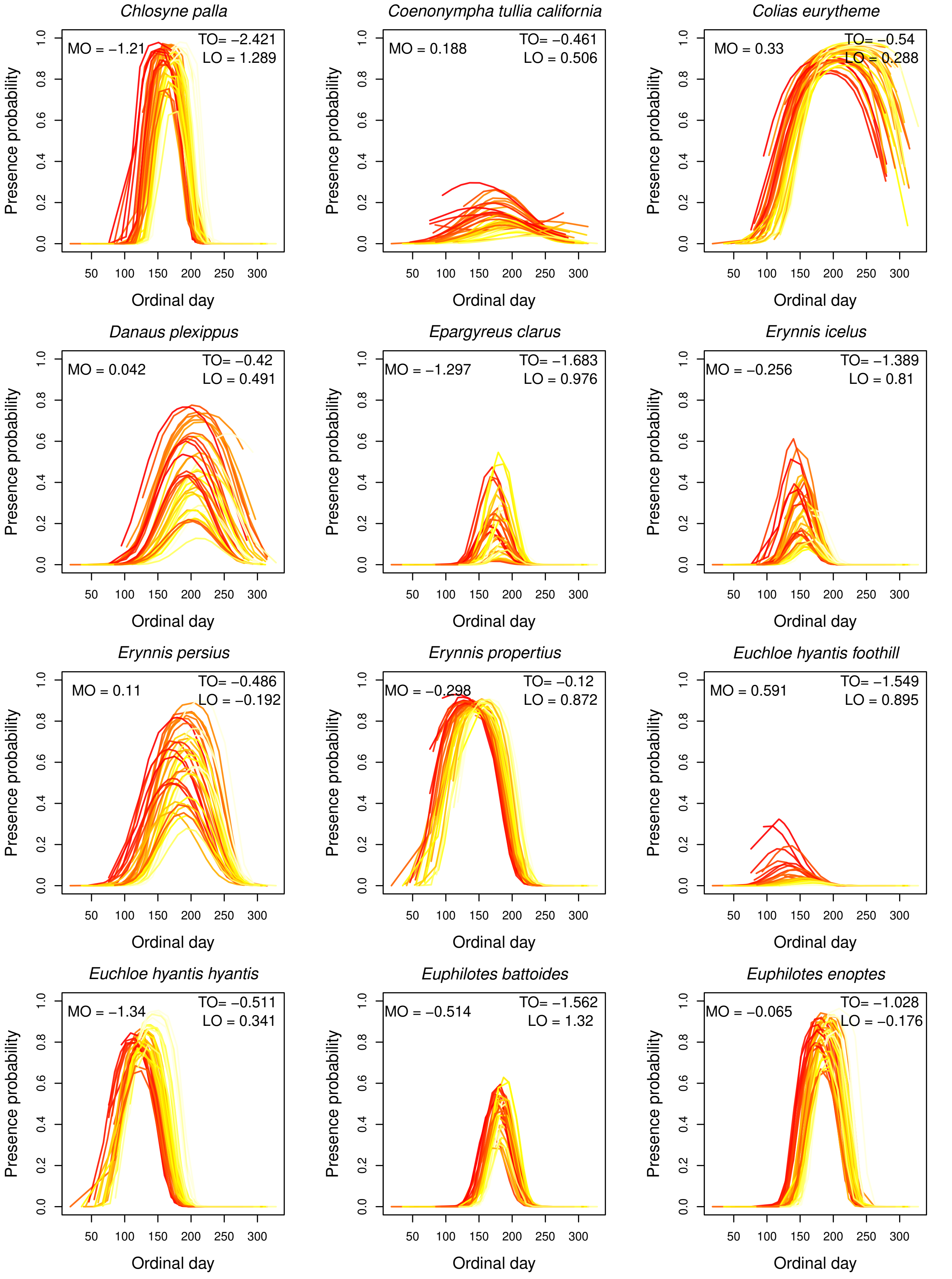


**FIGURE A17** Estimates of interannual variation in flight periods of species at Lang Crossing from our climate model. Each line represents the probability of occurrence on each day across a year, with colors indicating average spring maximum temperatures for that year (darker colors indicate higher spring maximum temperatures). Thus the effects of spring maximum temperature on MO, TO and LO for each species are shown [illegible]

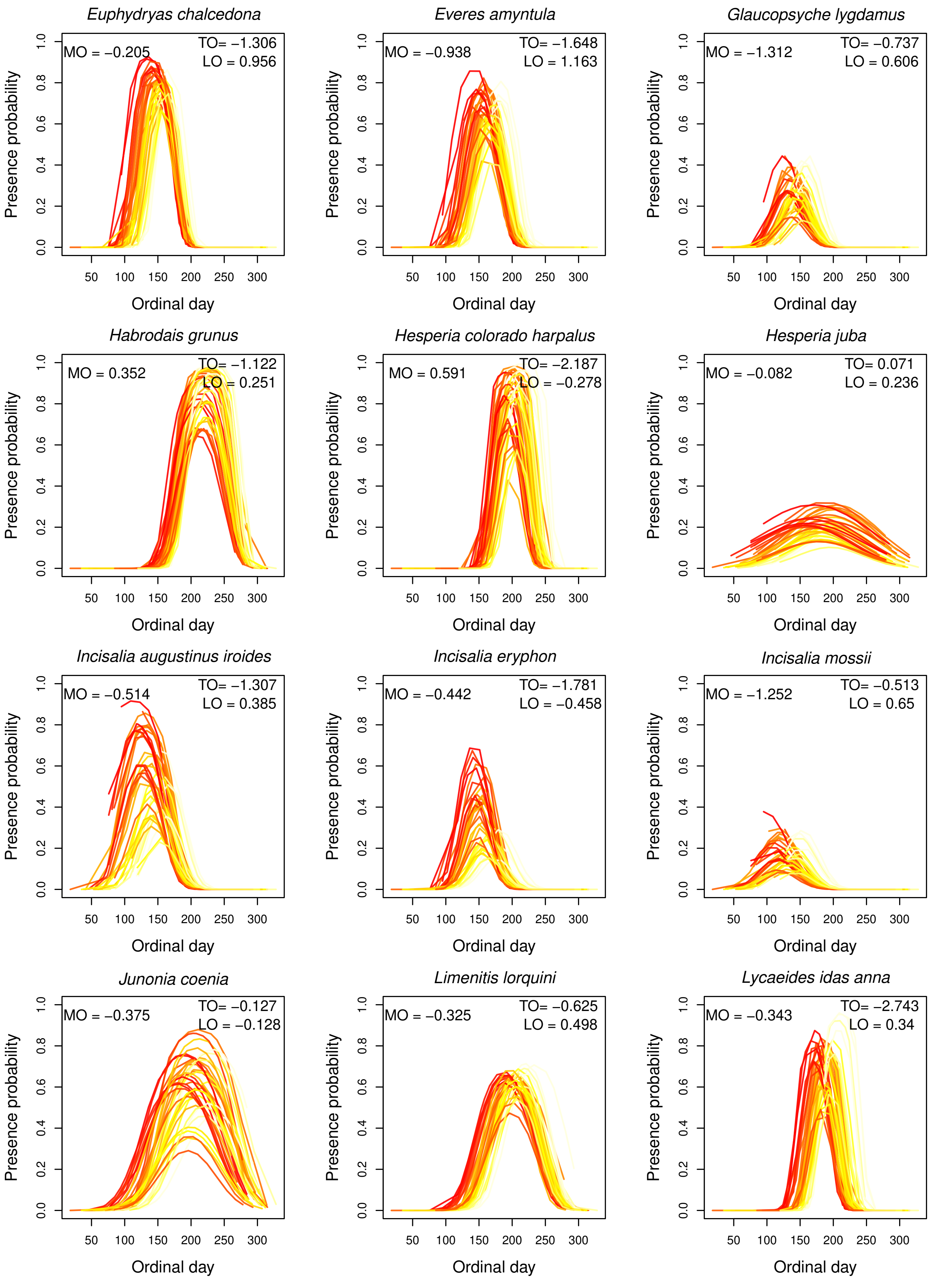


**FIGURE A18** Estimates of interannual variation in flight periods of species at Lang Crossing from our climate model. Each line represents the probability of occurrence on each day across a year, with colors indicating average spring maximum temperatures for that year (darker colors indicate higher spring maximum temperatures). Thus the effects of spring maximum temperature on MO, TO and LO for each species are shown [illegible]

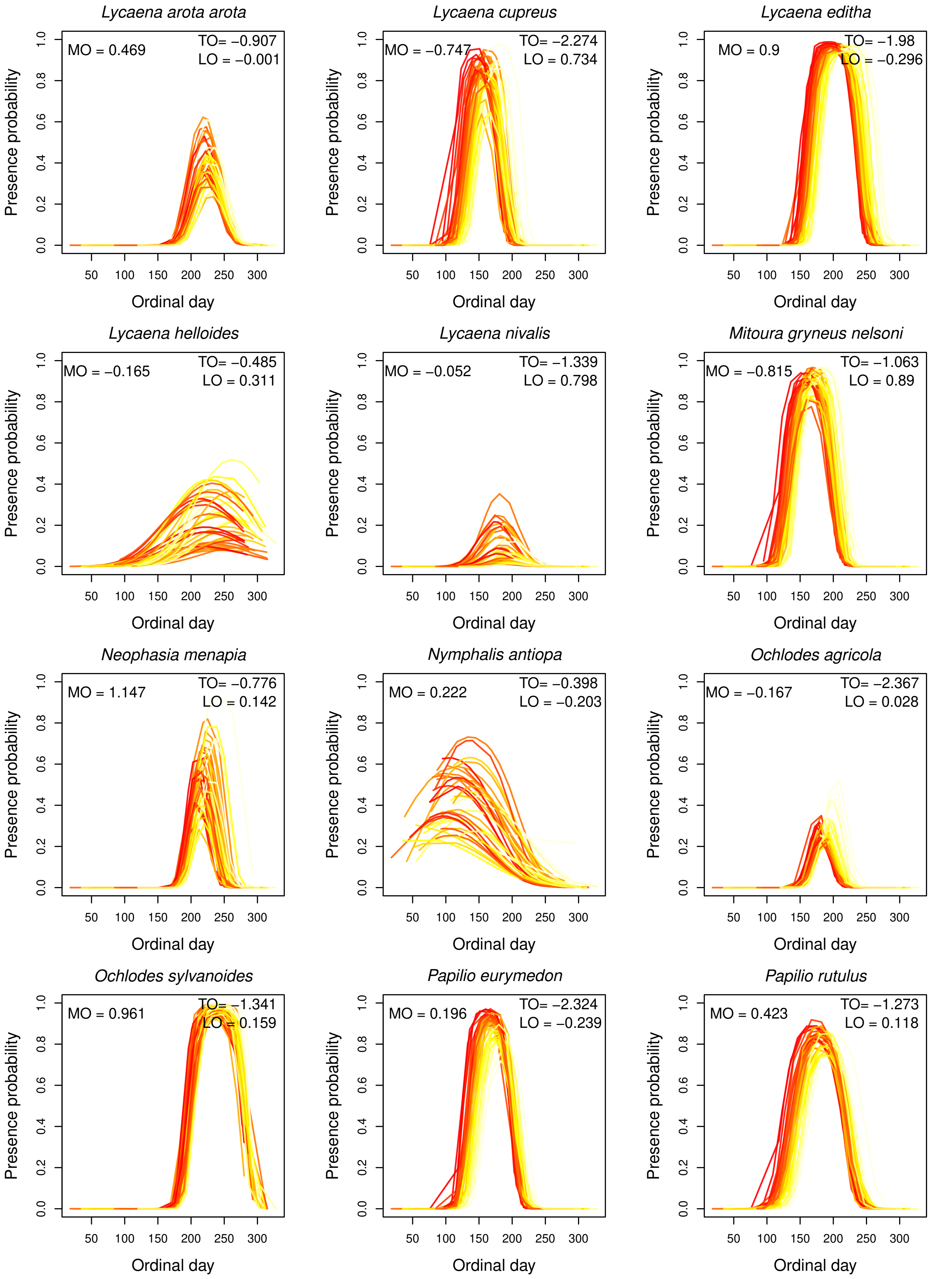


**FIGURE A19** Estimates of interannual variation in flight periods of species at Lang Crossing from our climate model. Each line represents the probability of occurrence on each day across a year, with colors indicating average spring maximum temperatures for that year (darker colors indicate higher spring maximum temperatures). Thus the effects of spring maximum temperature on MO, TO and LO for each species are shown [illegible]

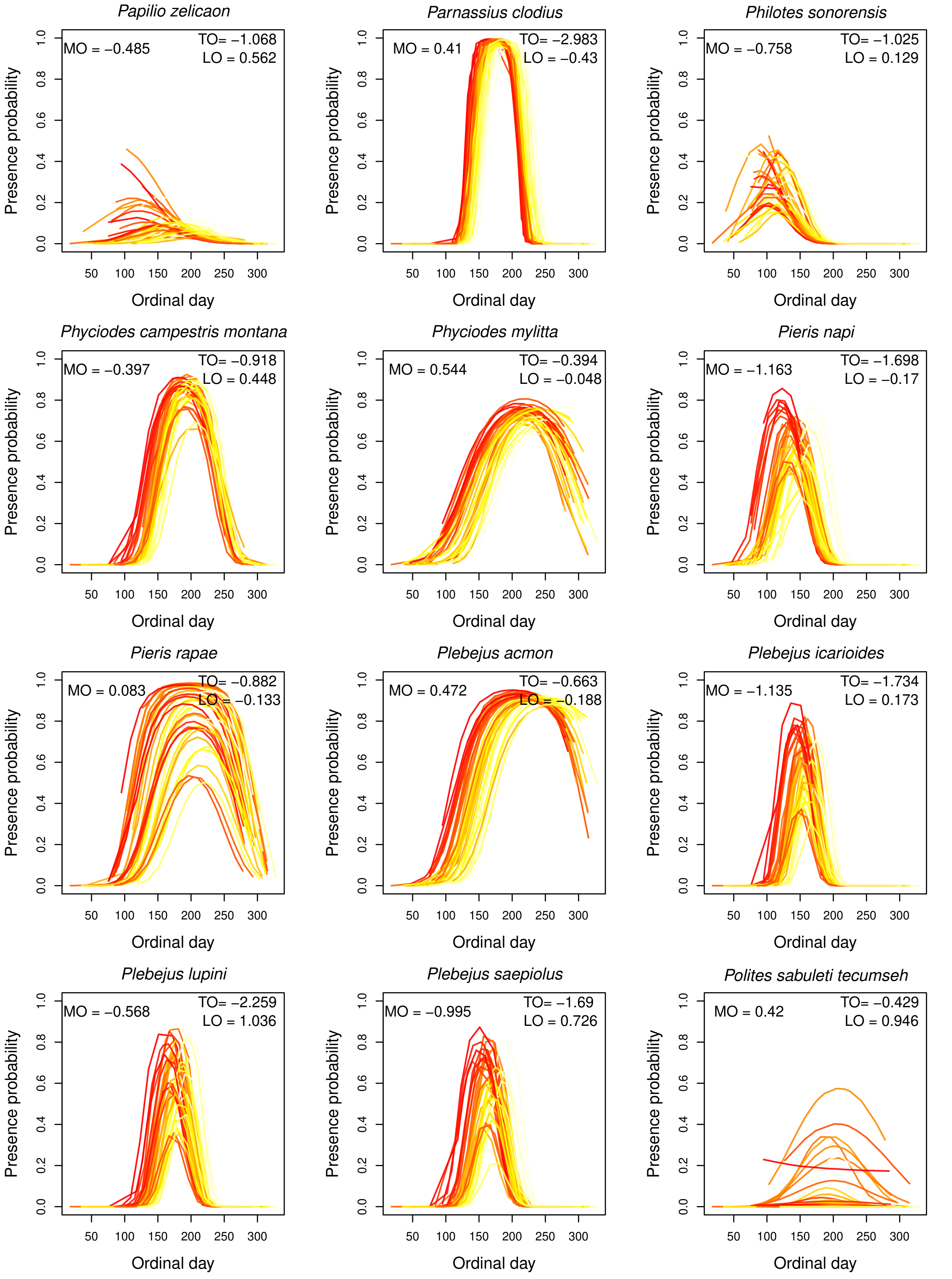


**FIGURE A20** Estimates of interannual variation in flight periods of species at Lang Crossing from our climate model. Each line represents the probability of occurrence on each day across a year, with colors indicating average spring maximum temperatures for that year (darker colors indicate higher spring maximum temperatures). Thus the effects of spring maximum temperature on MO, TO and LO for each species are shown [illegible]

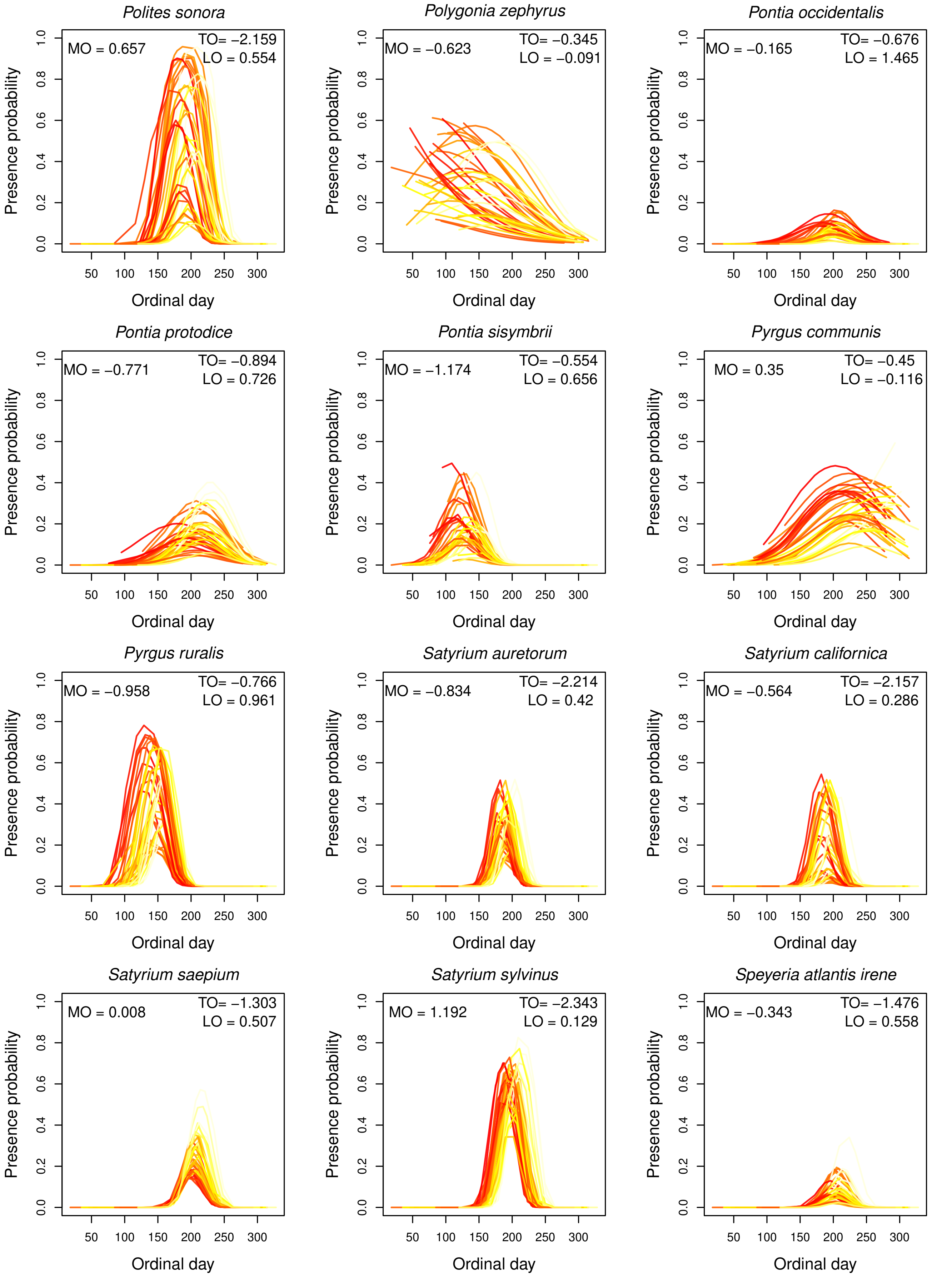


**FIGURE A21** Estimates of interannual variation in flight periods of species at Lang Crossing from our climate model. Each line represents the probability of occurrence on each day across a year, with colors indicating average spring maximum temperatures for that year (darker colors indicate higher spring maximum temperatures). Thus the effects of spring maximum temperature on MO, TO and LO for each species are shown [illegible]

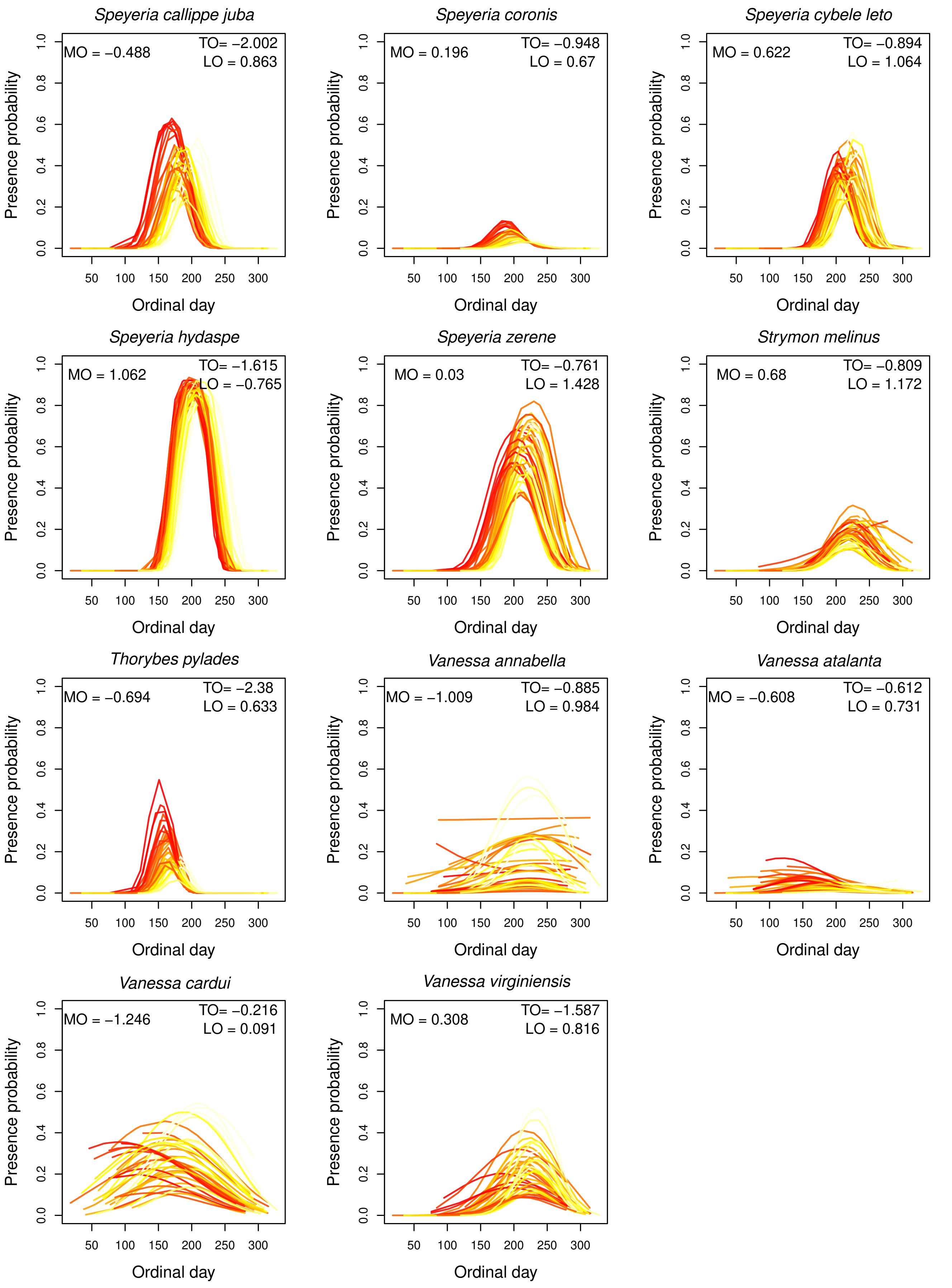


**FIGURE A22** Estimates of interannual variation in flight periods of species at Lang Crossing from our climate model. Each line represents the probability of occurrence on each day across a year, with colors indicating average spring maximum temperatures for that year (darker colors indicate higher spring maximum temperatures). Thus the effects of spring maximum temperature on MO, TO and LO for each species are shown [illegible]

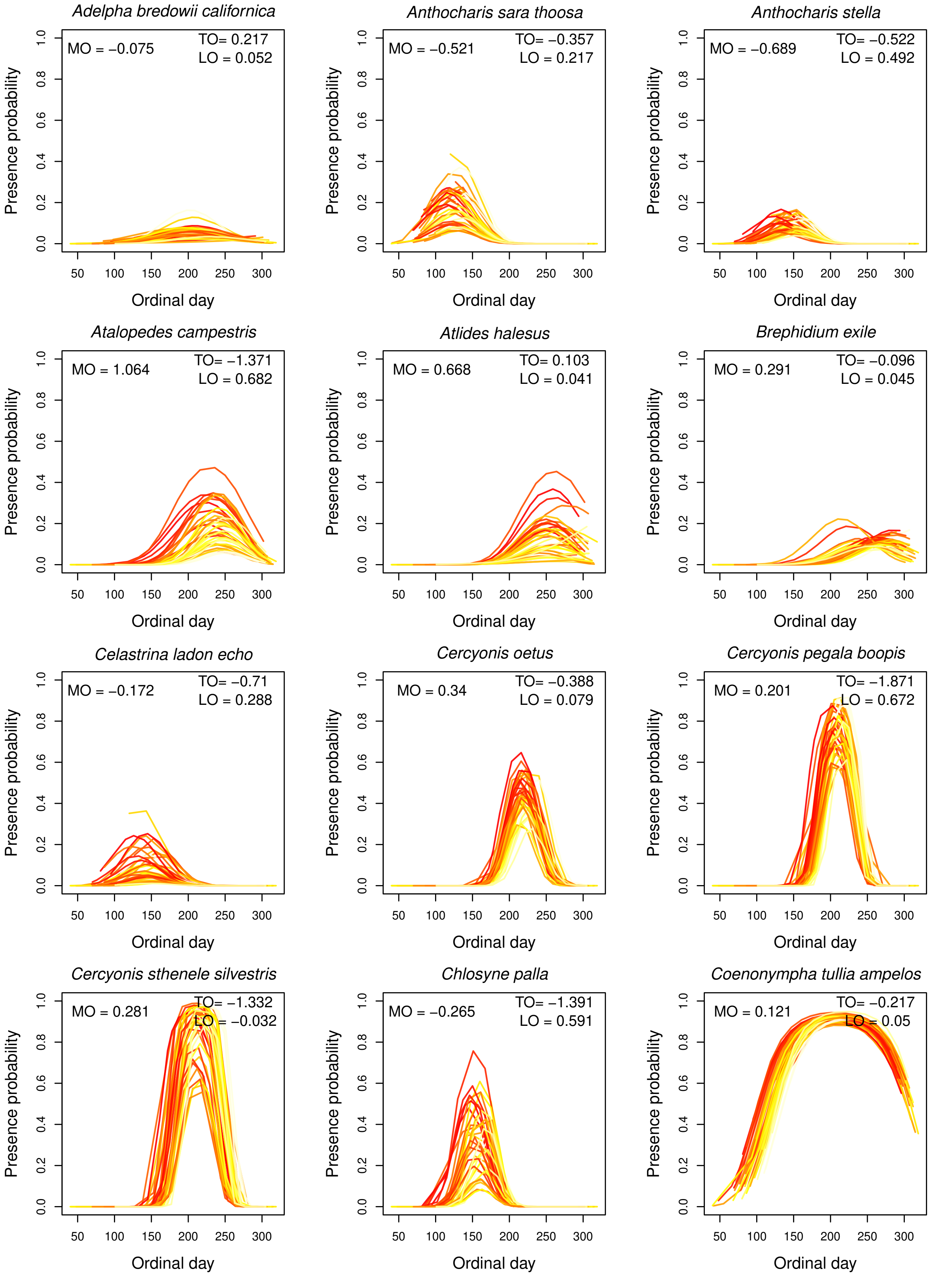


**FIGURE A23** Estimates of interannual variation in flight periods of species at Sierra Valley from our climate model. Each line represents the probability of occurrence on each day across a year, with colors indicating average spring maximum temperatures for that year (darker colors indicate higher spring maximum temperatures). Thus the effects of spring maximum temperature on MO, TO and LO for each species are shown [illegible]

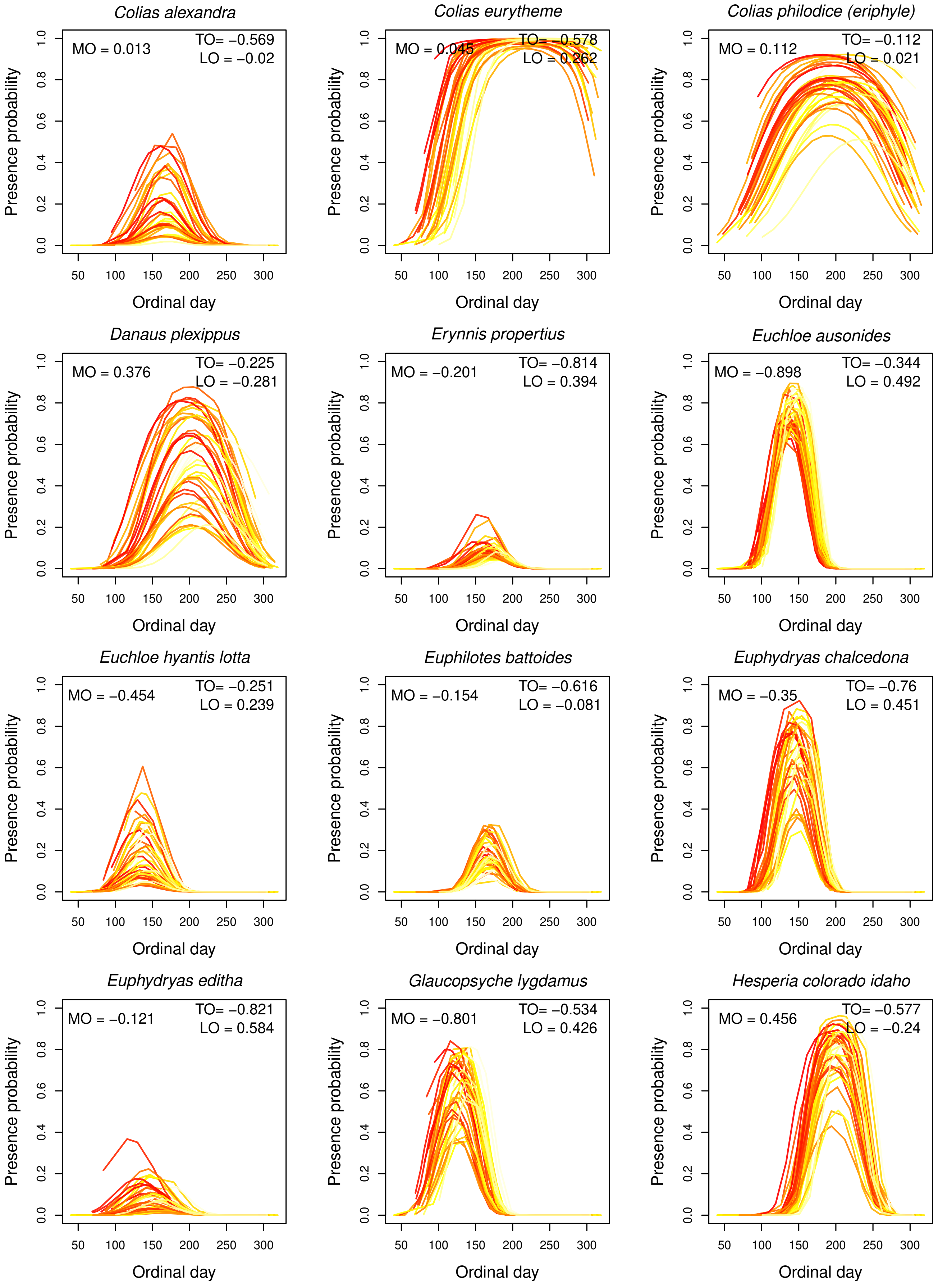


**FIGURE A24** Estimates of interannual variation in flight periods of species at Sierra Valley from our climate model. Each line represents the probability of occurrence on each day across a year, with colors indicating average spring maximum temperatures for that year (darker colors indicate higher spring maximum temperatures). Thus the effects of spring maximum temperature on MO, TO and LO for each species are shown [illegible]

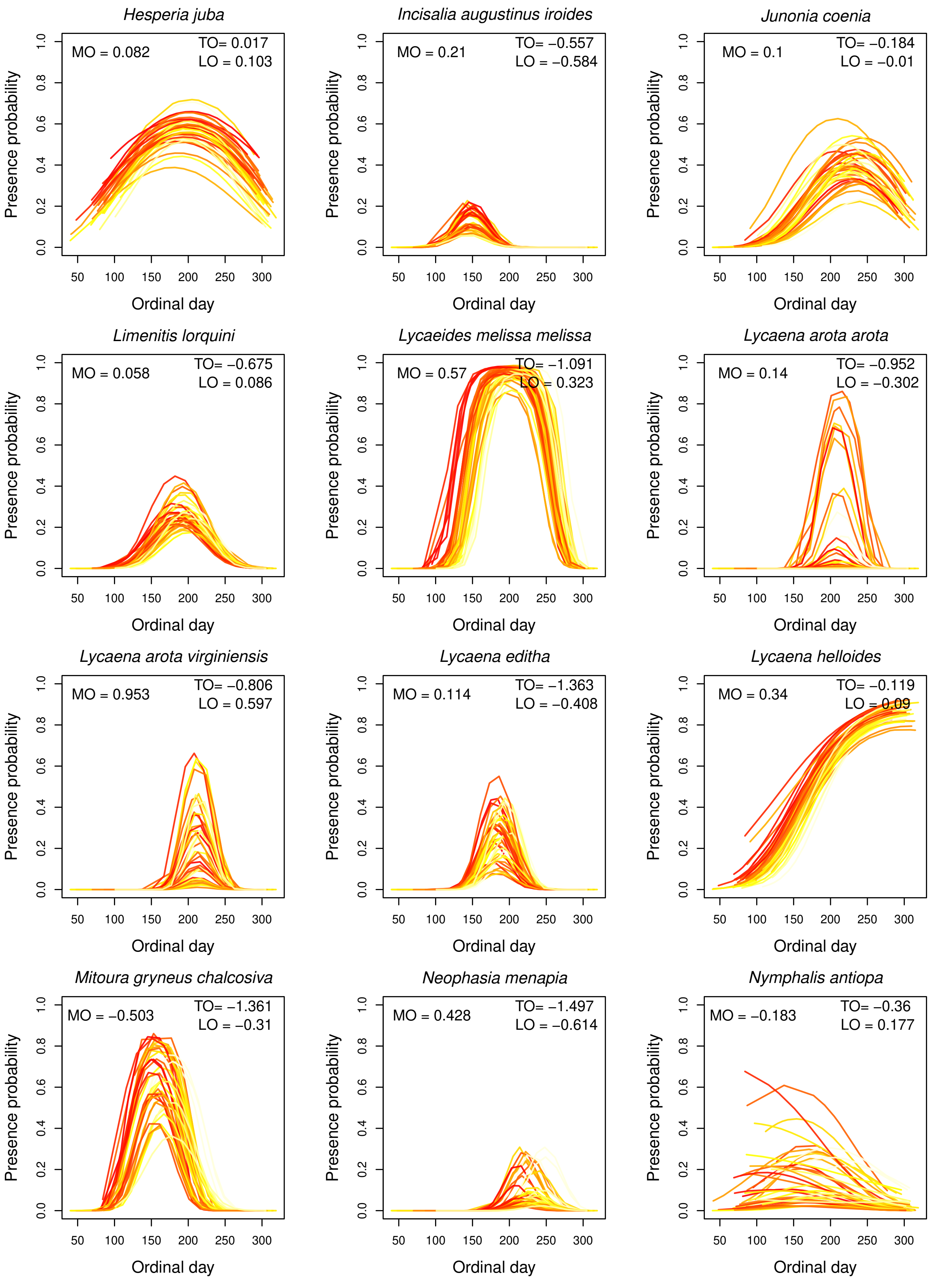


**FIGURE A25** Estimates of interannual variation in flight periods of species at Sierra Valley from our climate model. Each line represents the probability of occurrence on each day across a year, with colors indicating average spring maximum temperatures for that year (darker colors indicate higher spring maximum temperatures). Thus the effects of spring maximum temperature on MO, TO and LO for each species are shown [illegible]

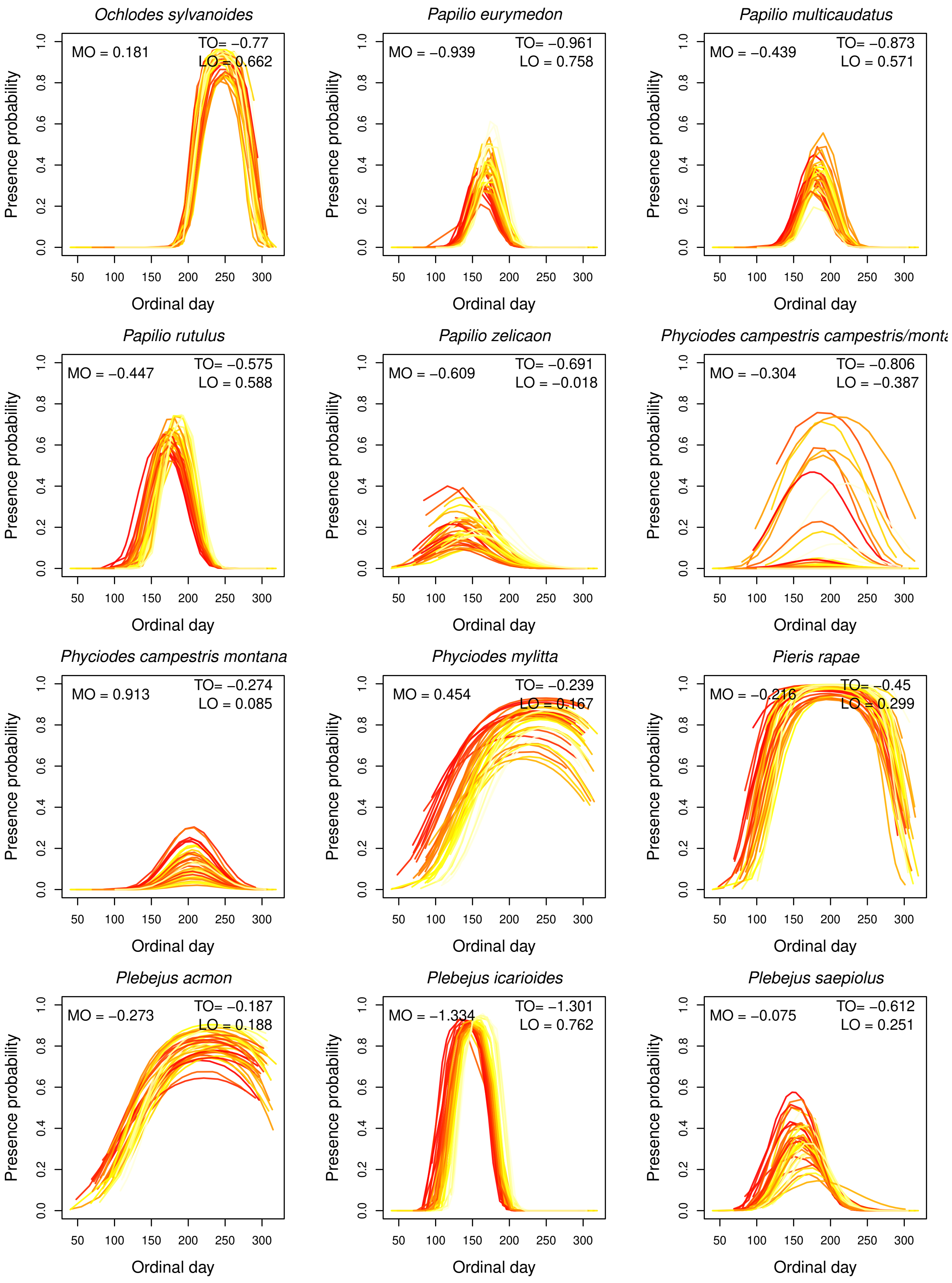


**FIGURE A26** Estimates of interannual variation in flight periods of species at Sierra Valley from our climate model. Each line represents the probability of occurrence on each day across a year, with colors indicating average spring maximum temperatures for that year (darker colors indicate higher spring maximum temperatures). Thus the effects of spring maximum temperature on MO, TO and LO for each species are shown [illegible]

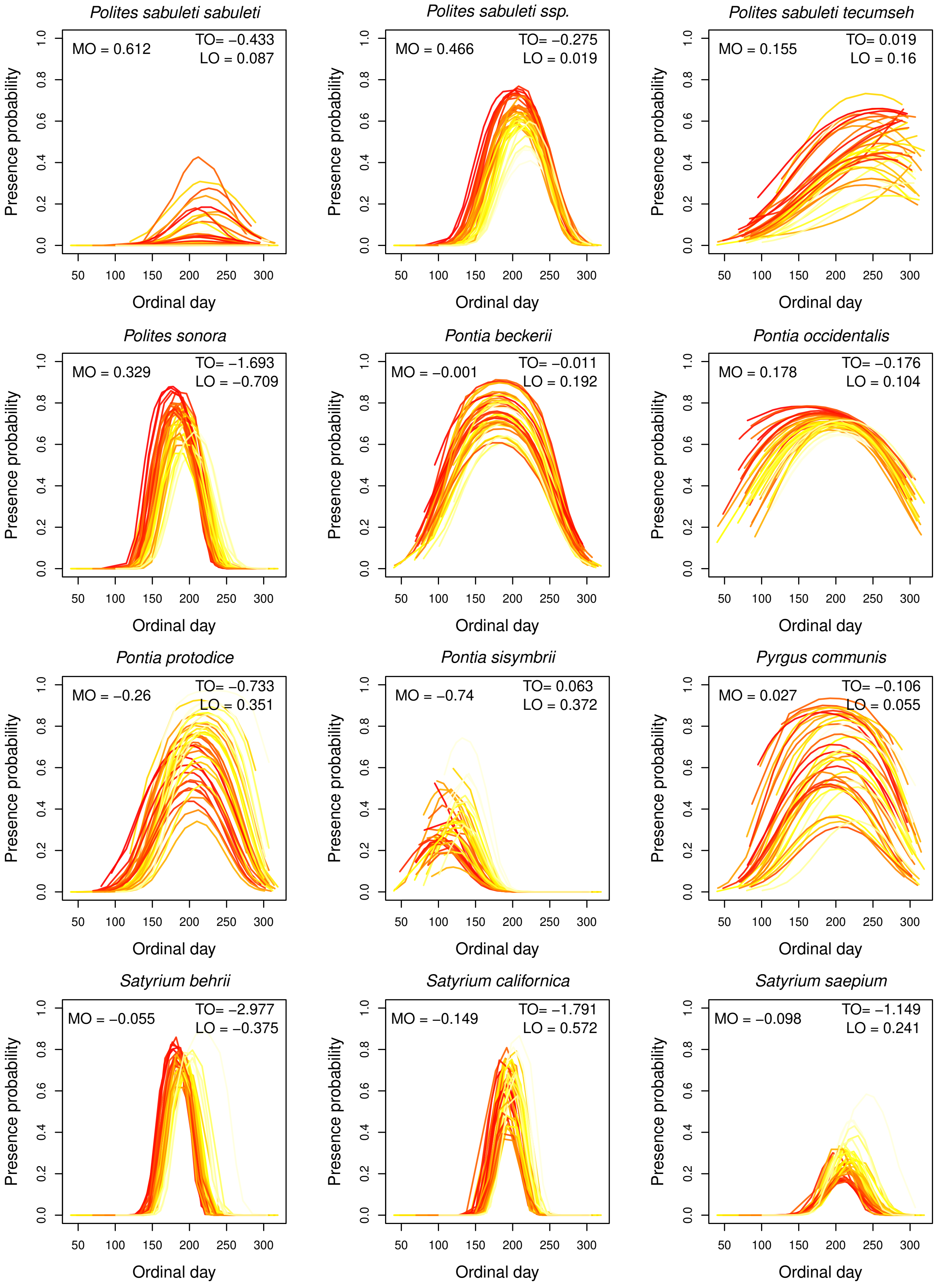


**FIGURE A27** Estimates of interannual variation in flight periods of species at Sierra Valley from our climate model. Each line represents the probability of occurrence on each day across a year, with colors indicating average spring maximum temperatures for that year (darker colors indicate higher spring maximum temperatures). Thus the effects of spring maximum temperature on MO, TO and LO for each species are shown [illegible]

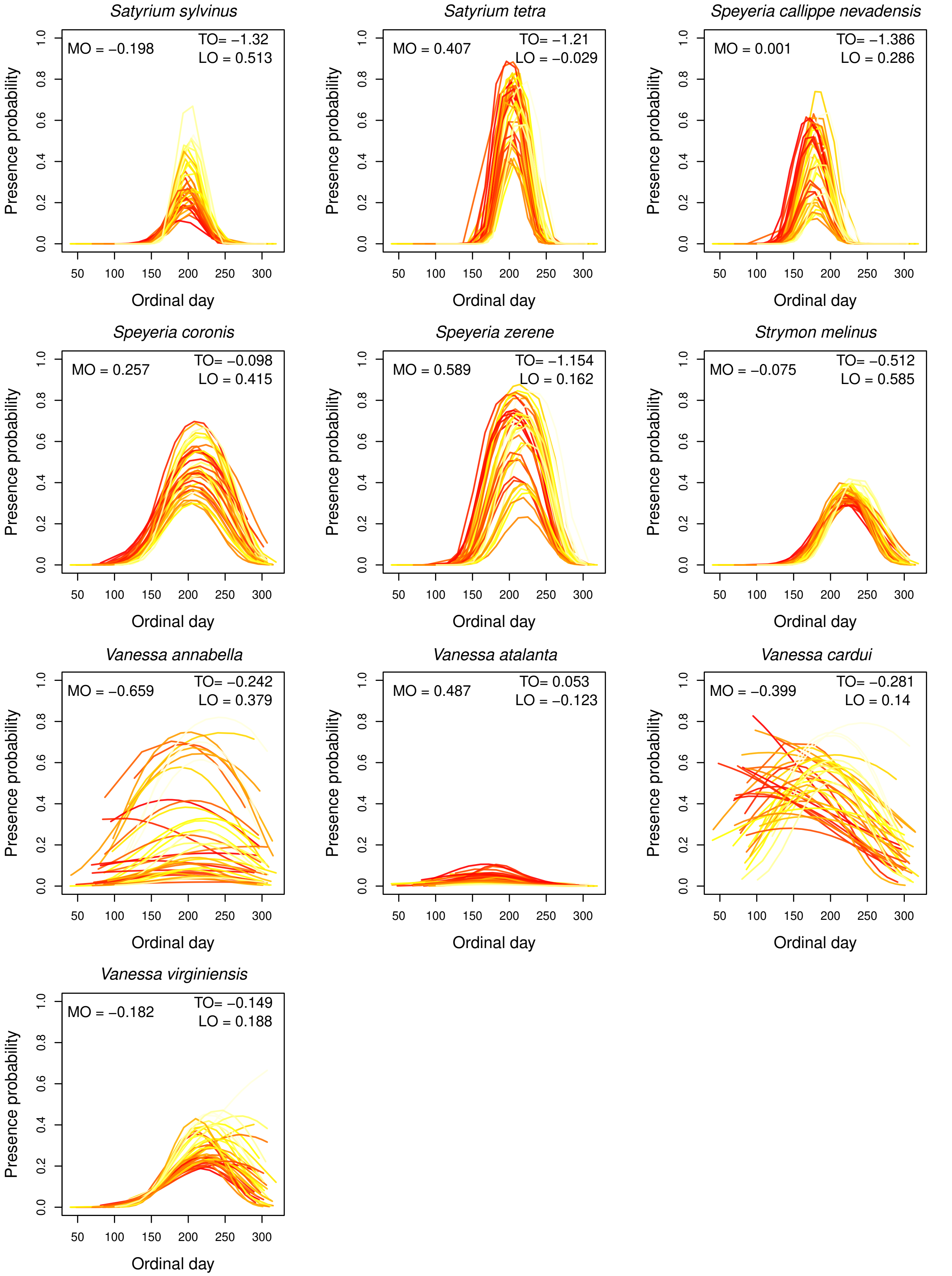


**FIGURE A28** Estimates of interannual variation in flight periods of species at Sierra Valley from our climate model. Each line represents the probability of occurrence on each day across a year, with colors indicating average spring maximum temperatures for that year (darker colors indicate higher spring maximum temperatures). Thus the effects of spring maximum temperature on MO, TO and LO for each species are shown in each plot.

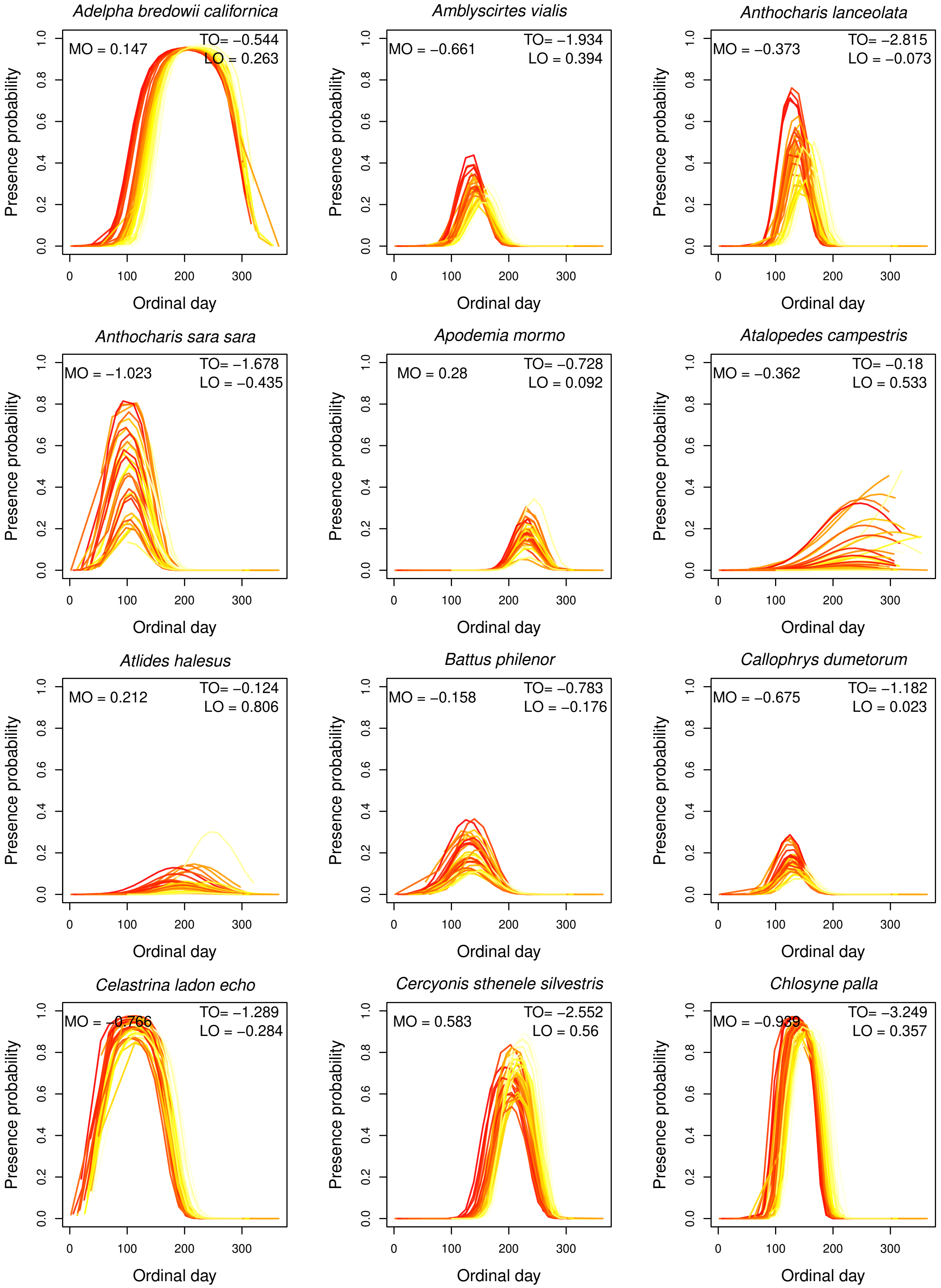


**FIGURE A29** Estimates of interannual variation in flight periods of species at Washington from our climate model. Each line represents the probability of occurrence on each day across a year, with colors indicating average spring maximum temperatures for that year (darker colors indicate higher spring maximum temperatures). Thus the effects of spring maximum temperature on MO, TO and LO for each species are shown [illegible]

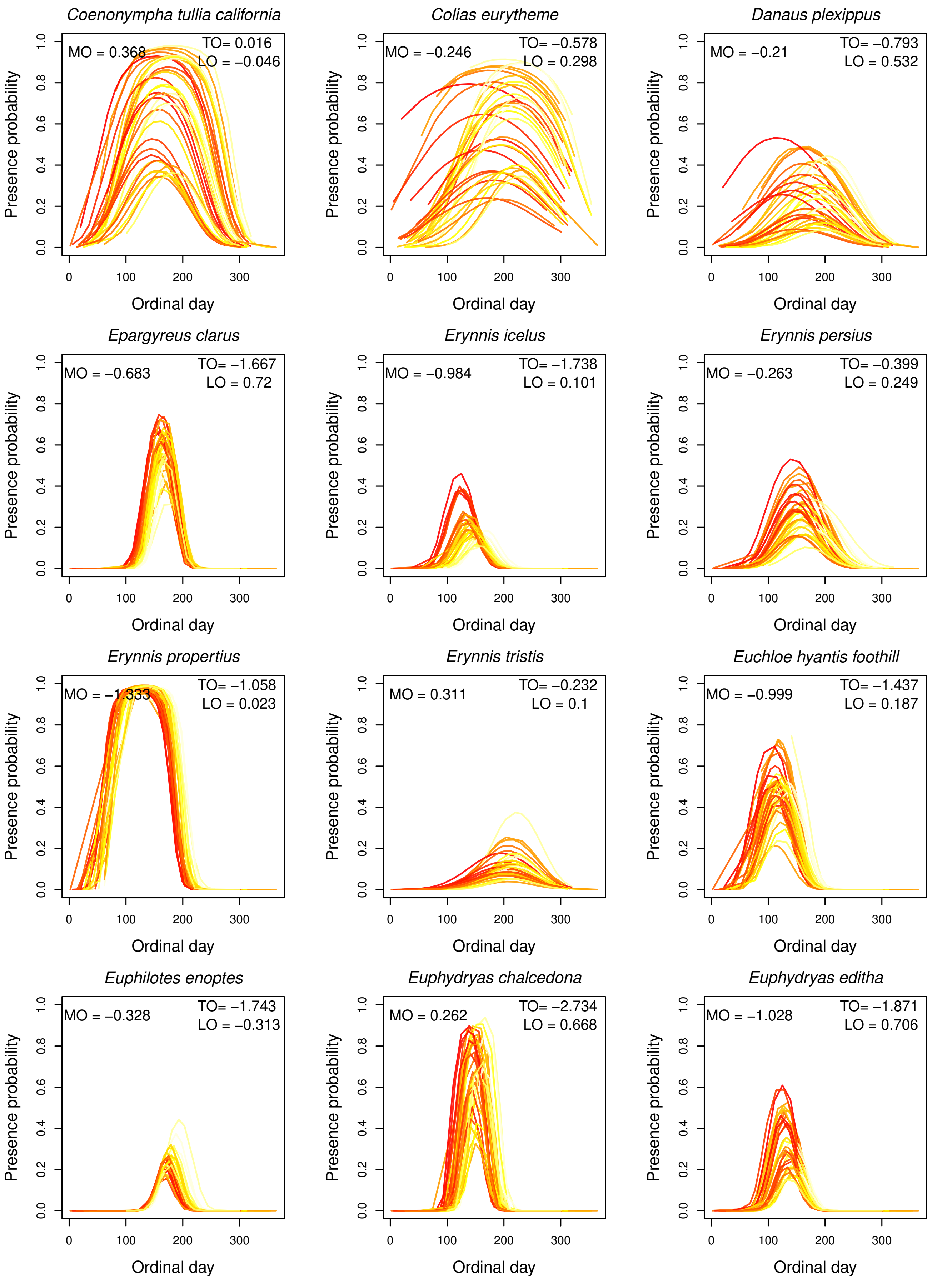


**FIGURE A30** Estimates of interannual variation in flight periods of species at Washington from our climate model. Each line represents the probability of occurrence on each day across a year, with colors indicating average spring maximum temperatures for that year (darker colors indicate higher spring maximum temperatures). Thus the effects of spring maximum temperature on MO, TO and LO for each species are shown [illegible]

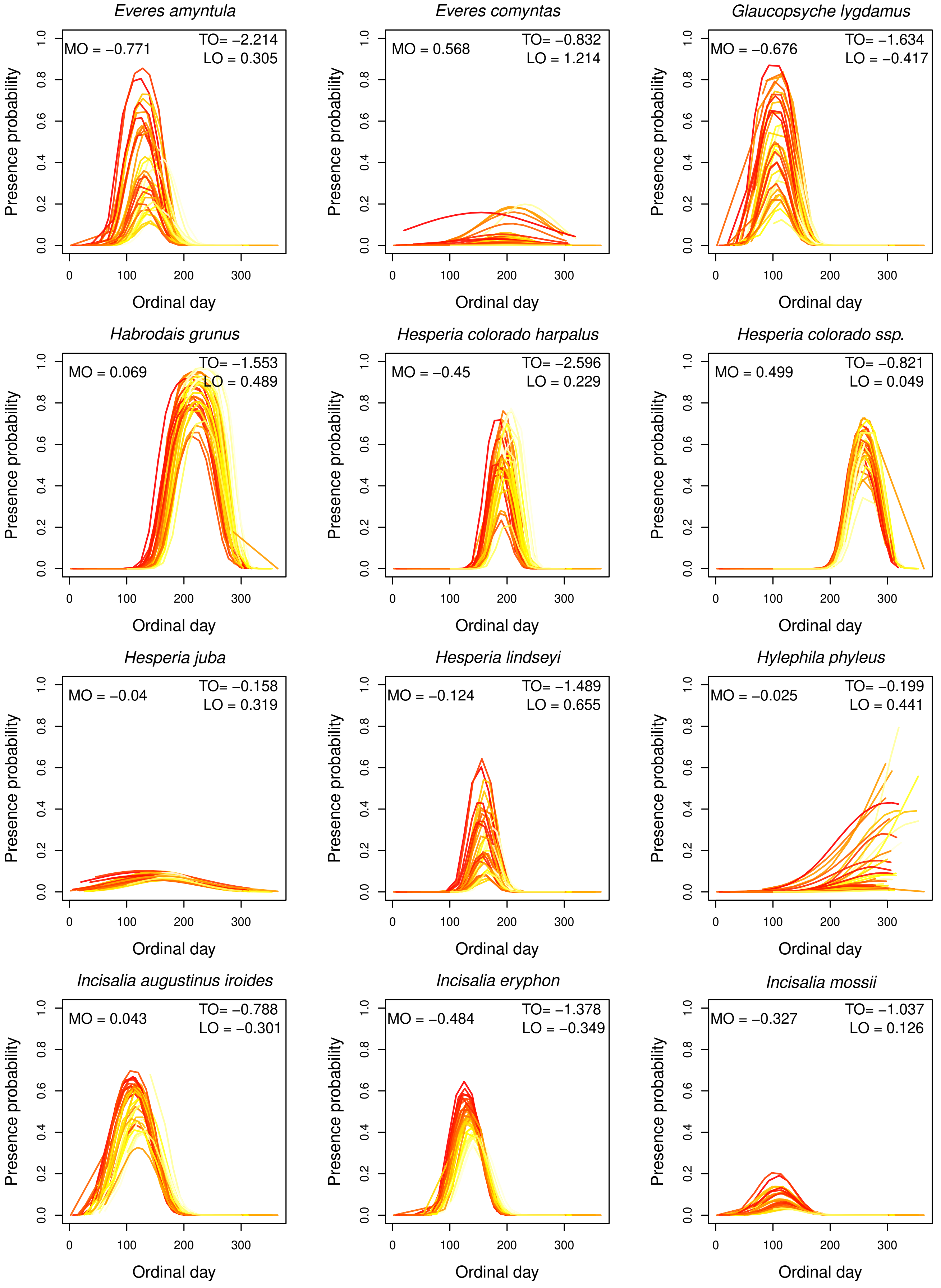


**FIGURE A31** Estimates of interannual variation in flight periods of species at Washington from our climate model. Each line represents the probability of occurrence on each day across a year, with colors indicating average spring maximum temperatures for that year (darker colors indicate higher spring maximum temperatures). Thus the effects of spring maximum temperature on MO, TO and LO for each species are shown [illegible]

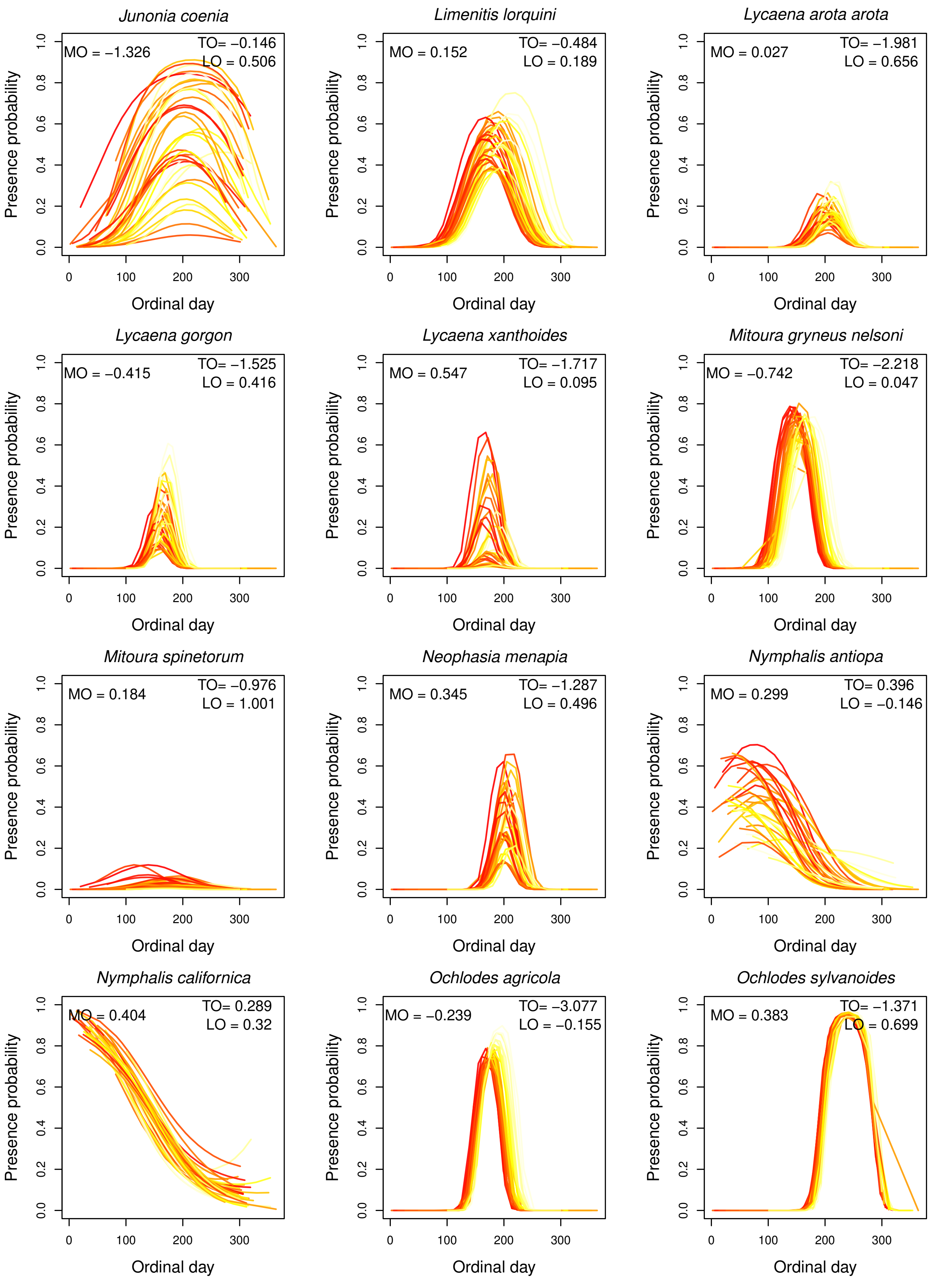


**FIGURE A32** Estimates of interannual variation in flight periods of species at Washington from our climate model. Each line represents the probability of occurrence on each day across a year, with colors indicating average spring maximum temperatures for that year (darker colors indicate higher spring maximum temperatures). Thus the effects of spring maximum temperature on MO, TO and LO for each species are shown [illegible]

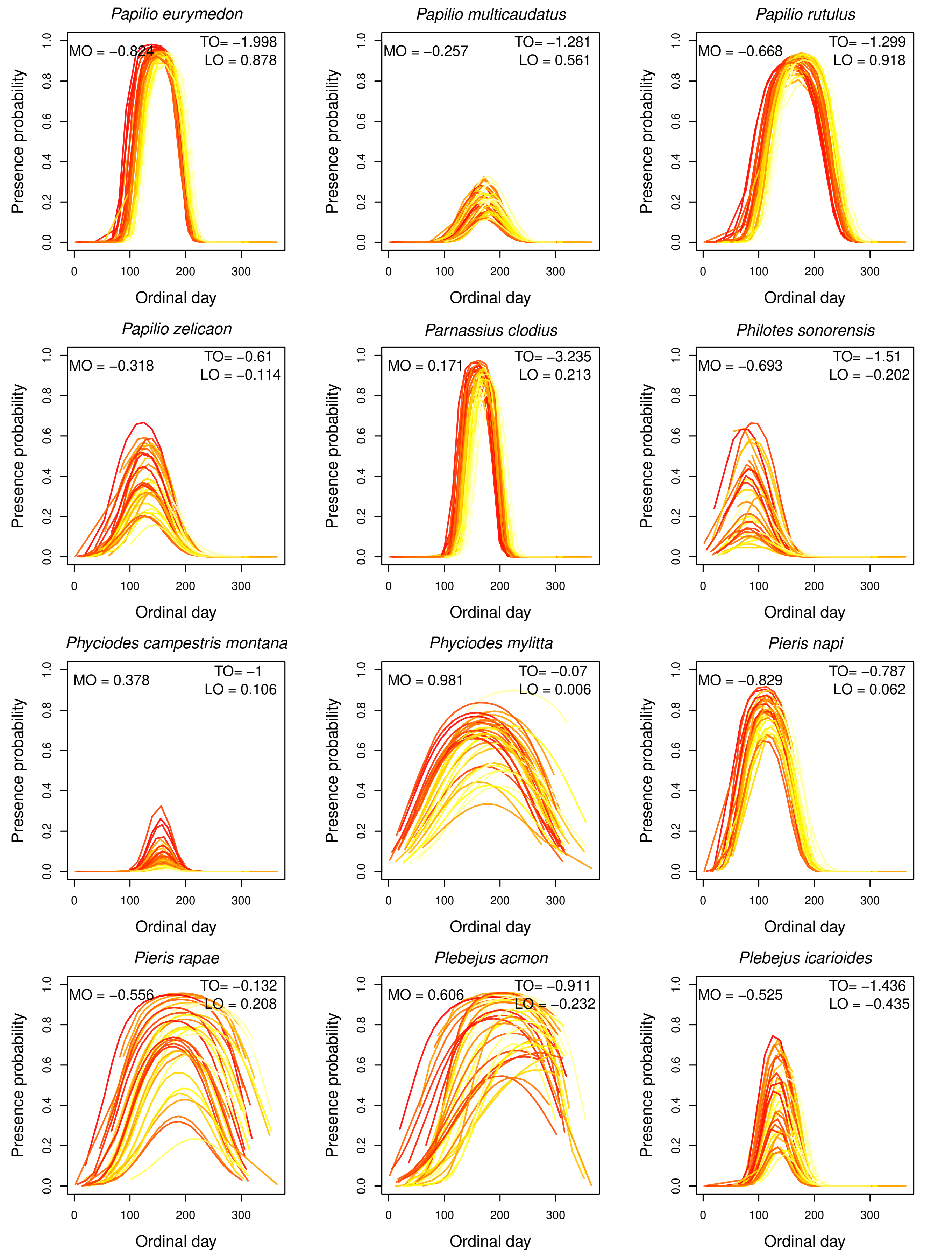


**FIGURE A33** Estimates of interannual variation in flight periods of species at Washington from our climate model. Each line represents the probability of occurrence on each day across a year, with colors indicating average spring maximum temperatures for that year (darker colors indicate higher spring maximum temperatures). Thus the effects of spring maximum temperature on MO, TO and LO for each species are shown in each plot.

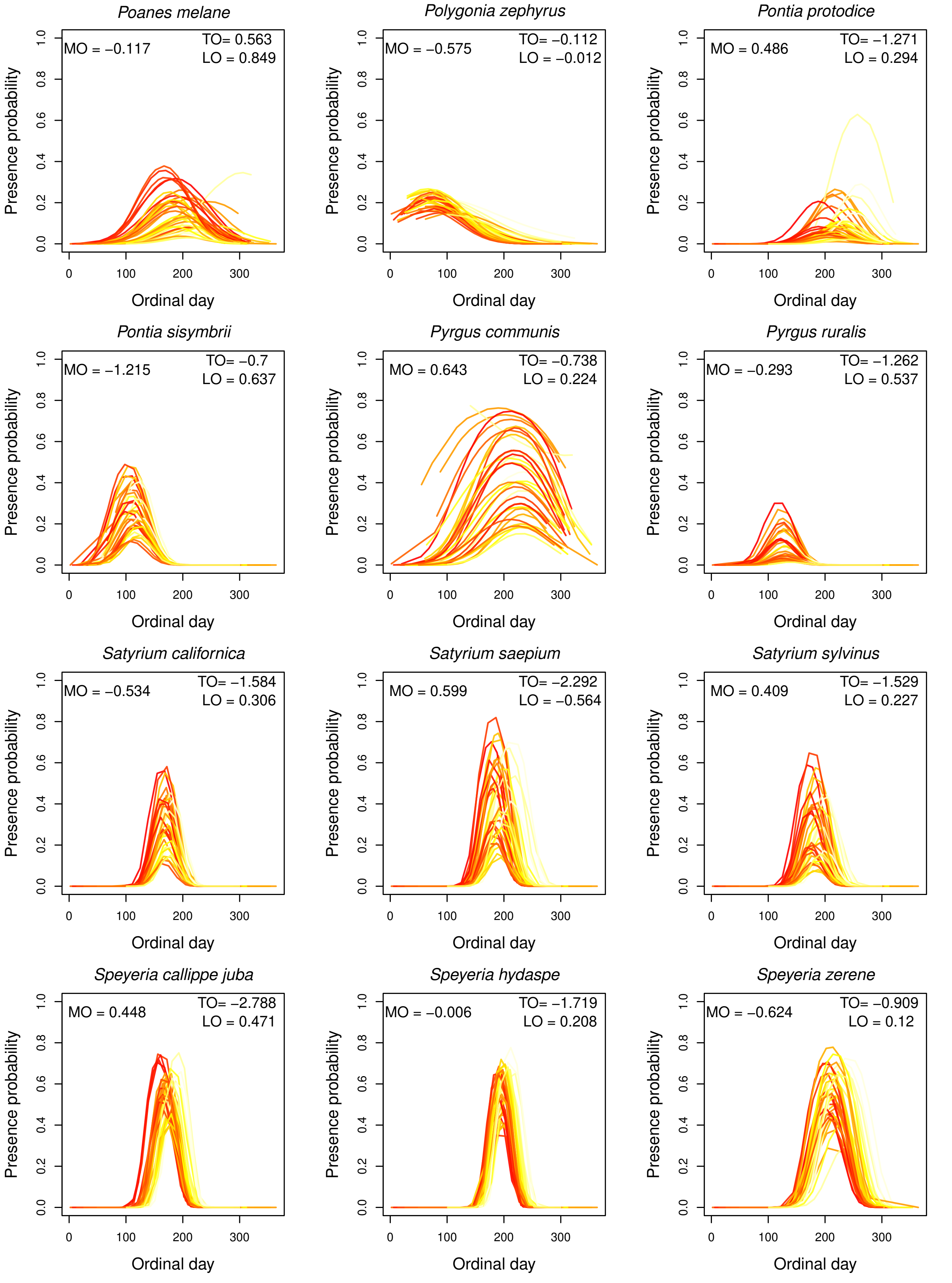


**FIGURE A34** Estimates of interannual variation in flight periods of species at Washington from our climate model. Each line represents the probability of occurrence on each day across a year, with colors indicating average spring maximum temperatures for that year (darker colors indicate higher spring maximum temperatures). Thus the effects of spring maximum temperature on MO, TO and LO for each species are shown [illegible]

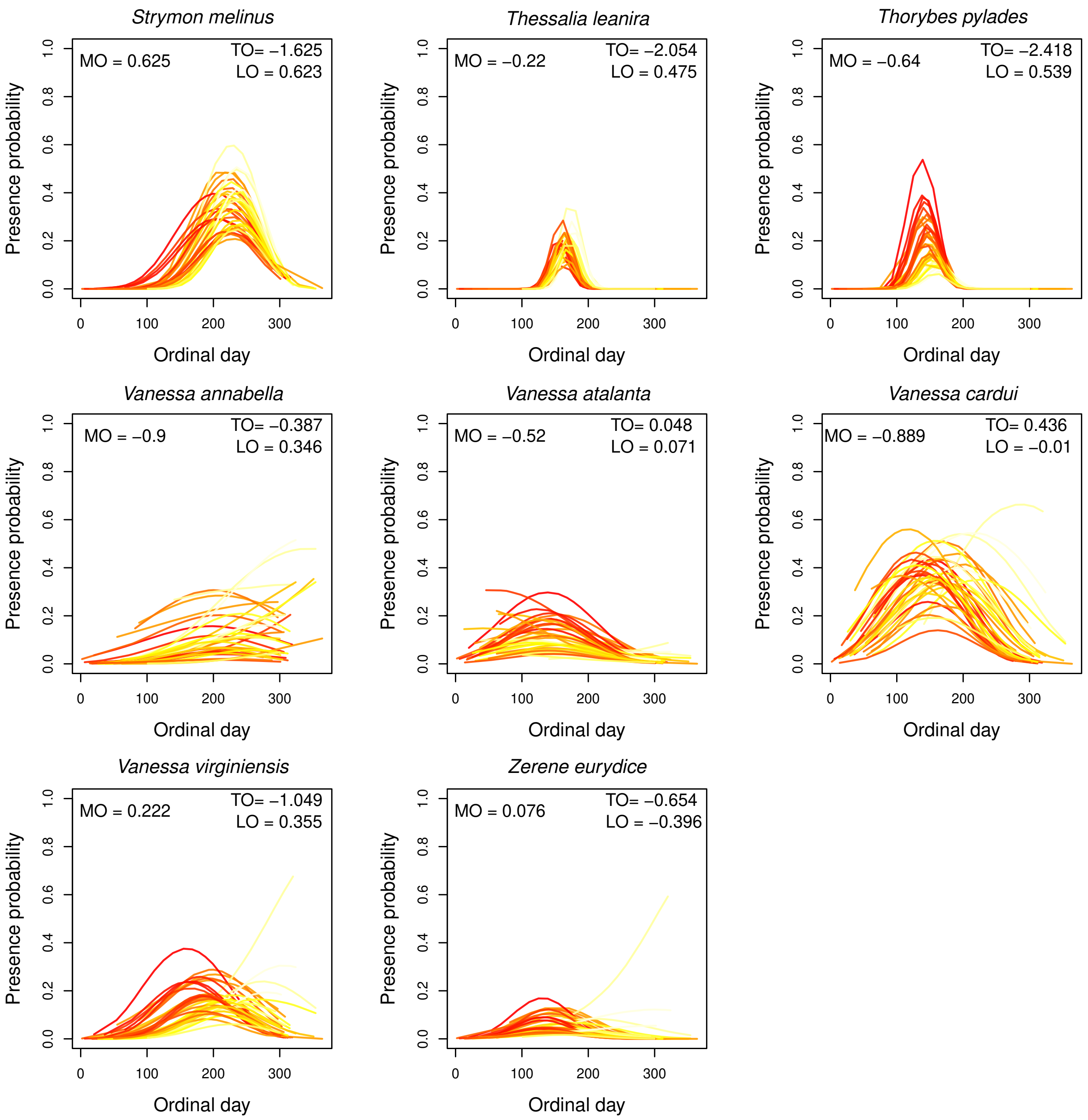


**FIGURE A35** Estimates of interannual variation in flight periods of species at Washington from our climate model. Each line represents the probability of occurrence on each day across a year, with colors indicating average spring maximum temperatures for that year (darker colors indicate higher spring maximum temperatures). Thus the effects of spring maximum temperature on MO, TO and LO for each species are shown [illegible]

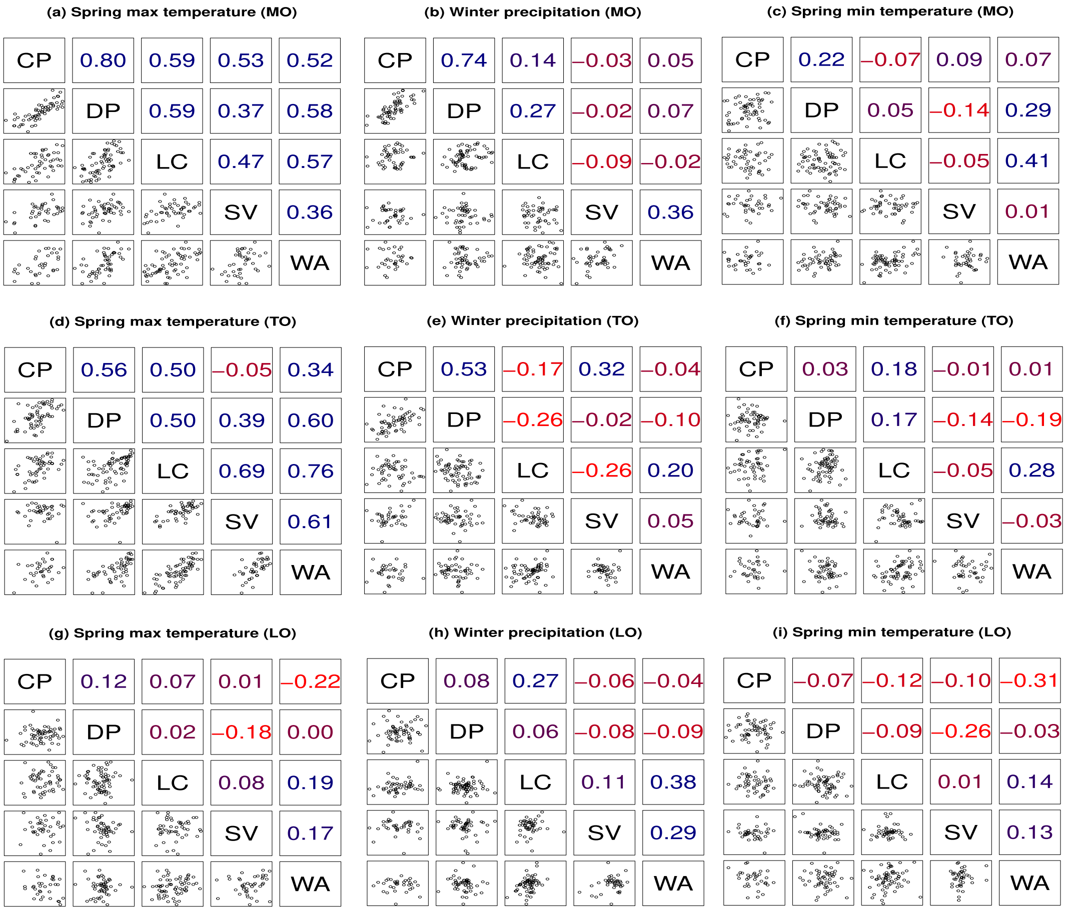


**FIGURE A36** Similarity in the effect of each climatic variable on butterfly occurrence distribution across sites. Results are shown for the effect of (a) spring maximum temperature on the mid-season occurrence (MO), (b) winter precipitation on the mid-season occurrence (MO), (c) spring minimum temperature on the mid-season occurrence (MO), (d) spring maximum temperature on the timing of occurrence (TO), (e) winter precipitation on the timing of occurrence (TO), (f) spring minimum temperature on the timing of occurrence (TO), (g) spring maximum temperature on length of occurrence (LO), (h) winter precipitation on length of occurrence (LO), (i) spring minimum temperature on length of occurrence (LO). In each panel, each point on the scatter plot represents a specific species present in both compared sites. The correlation coefficient indicates the similarity in the effect of each climatic variable on the flight period of species observed in both compared sites. The higher the correlation coefficient, the darker the color, with red representing negative correlation and blue representing positive correlation. A stronger correlation signifies a higher similarity in the effect of climate on species present at both sites.

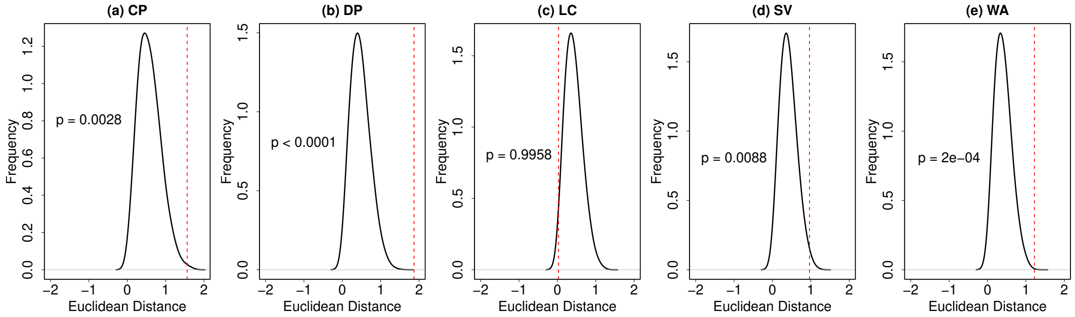


**FIGURE A37** Distribution of Euclidean distances from 10,000 permutations (black curve) testing whether the observed relationship between climate effects and different aspects of the flight period differs significantly between multivoltine and univoltine species at (a) Castle Peak, (b) Donner Pass, (c) Lang Crossing, (d) Sierra Valley, and (e) Washington. The red dashed line represents the observed Euclidean distance. P-values indicate whether the observed difference is statistically significant.
